# Age-Related Changes to Olivocochlear Efferent Neurons in Mice with Pathological Hearing

**DOI:** 10.64898/2025.12.27.696605

**Authors:** Kirupa Suthakar, Hannah Douglas-Kinniburgh, David K. Ryugo

## Abstract

One of the fundamental features of age-related hearing loss (ARHL) is difficulty discriminating speech signals from background noise. In addition to protecting the ear from acoustic trauma, olivocochlear (OC) efferent neurons participate in signal discrimination by virtue of their inhibitory actions on auditory nerve firing. Given the time course of peripheral degeneration in ARHL, we sought to investigate the central degeneration of medial (MOC) and lateral (LOC) efferent neurons in mutant mice that exhibit genetic hearing loss or deafness at different ages. Tests of cochlear function were combined with anatomical and morphological quantification of changes to somatic number, morphology, and location of OC neurons. Neuronal tract tracing methods were employed to label OC neurons in 1-, 3-, and 6-month-old CBA/CaH mice with normal hearing; DBA/2J mice with progressive, high frequency hearing loss; and homozygous Shaker2 mice with congenital deafness. Deaf Shaker2-/- animals exhibited age-related atrophy and loss of MOCs, with contralateral MOCs more affected than ipsilateral MOCs, while LOCs were largely unaffected. No such OC degeneration was observed in DBA/2J mice, even after progressive elevation of low frequency auditory brainstem response (ABR) thresholds and distortion product otoacoustic emissions (DPOAE) thresholds. Thus, OC efferent neurons can appear morphologically ‘normal’ in the complete absence of acoustic input in early life (as in deaf Shaker2-/- animals) and that the retention of these neurons is not affected by late onset high-frequency hearing loss observed in DBA/2J animals. Differential patterns of MOC neuron degeneration may affect functional plasticity of auditory brainstem feedback circuitry in ARHL.

## Introduction

Within the auditory system, there are two mechanisms of efferent modulation: the middle ear muscle reflex (MEMR) and auditory efferent neurons, comprising the classical olivocochlear (OC) neurons and the recently identified dorsal efferent neurons (DEs; Suthakar and Ryugo 2021). Both MEMR and OC efferent systems are sound-activated negative feedback arcs differing in their respective peripheral innervation patterns to the muscles of the middle ear (Borg 1973) or cells of the inner ear (Guinan 2011).

In most mammals, the OC neurons can be divided into a medial (MOC) and lateral (LOC) group based on their somatic locations in the superior olivary complex (SOC). MOCs and LOCs can also be morphologically differentiated by their immunohistochemical profiles, axonal myelination, and peripheral projection patterns (Warr, Boche et al. 1997). MOC somata are large and stellate, located in periolivary regions of the superior olivary complex (SOC; Osen, Mugnaini et al. 1984, Warr 1992, Brown and Levine 2008). LOCs are disc shaped, located in the lateral superior olive, predominantly ipsilateral to the cochlea they innervate. Recent data have demonstrated that LOCs are molecularly diverse (Frank, Sitko et al. 2023) and can adopt a dopaminergic identity in response to traumatic noise exposure (Wu, Yi et al. 2020, Sitko, Frank et al. 2025).

In mouse, MOCs are located bilaterally in the ventral nucleus of the trapezoid body (VNTB), which occupies the space lateral to the medial nucleus of the trapezoid body (MNTB), medial to the lateral superior olive (LSO), and ventral to the superior paraolivary nucleus. At the rostral extent of the SOC, the VNTB migrates dorsolaterally to form the rostral periolivary nucleus (RPO). The RPO continues past the rostral extent of the MNTB and is continuous with the ventral nucleus of the lateral lemniscus (VNLL), where DEs are found. MOCs can be subdivided based on their somatic location relative to the cochlea they innervate (contralateral MOCs or ipsilateral MOCs), or relative to the cochlea from which their primary afferent input is received (contralateral, ipsilateral, or bilateral units; Warr 1992, Thompson and Schofield 2000). This situation ensures that the MOC pathway to either ear can incorporate afferent sound information from ***both*** ears.

The distribution of MOCs either ipsilateral or contralateral to the ear they innervate is an important consideration when assessing their functionality. The double-crossed nature of the MOC system ensures that each ear receives efferent innervation driven by sound input from the same side (via contralateral MOCs) and the opposite side (via ipsilateral MOCs). Considering that sound localization processing is performed by adjacent SOC nuclei, this situation places OC neurons in an ideal position to receive information necessary to compute relative differences required for sound source location when background noise is high. There is evidence to support such a projection pattern. For example, in addition to axonal input to glycinergic VNTB neurons, axons of principal medial nucleus of the trapezoid body (MNTB) cells which are involved in calculating interaural time differences form collaterals with non-glycinergic cells which are presumably the cholinergic MOC neurons (Spangler, Warr et al. 1985, Kuwabara and Zook 1991, Albrecht, Dondzillo et al. 2014). Indeed, recent in-vitro whole cell patch clamp electrophysiology supports a direct functional GABAergic synapse from MNTB to MOC neurons (Torres Cadenas, Fischl et al. 2020, Torres Cadenas, Cheng et al. 2022).

Disruption of sensory input is known to cause central nervous system disorganization (Hubel and Wiesel 1970, Van der Loos and Woolsey 1973, Benson, Ryugo et al. 1984). In the auditory system, peripheral axotomy via cochlear ablation has been used to investigate effects of deafness in the brain (Powell and Cowan 1962, Kane 1974b, Nordeen, Killackey et al. 1983, Kitzes and Semple 1985, Moore 1985). Specific to OC efferent neurons, cochleotomies in adult rats result in an age-dependent reduction in the number of LOCs, but not MOCs (Kraus and Illing 2005). This differential survival pattern may be explained by the retention of potential non-auditory inputs to MOCs that would not be affected by unilateral cochlear ablation. One significant complication to surgical ablation is possible involvement of other auditory tissue (spiral ganglion) and disruption of the vascular supply to nearby brainstem structures.

Another model used to study the effects of deafness on OCs is congenital hypothyroidism. This condition affects organ of Corti development, resulting in hypertrophy of the tectorial membrane (Uziel, Gabrion et al. 1981) and OHC abnormalities (Prieto, Rueda et al. 1990), consequently causing elevated DPOAE and ABR thresholds (Li, Henley et al. 1999). Contrary to the situation following cochlear ablation, congenitally deaf hypothyroid rats exhibited normal neuronal counts, but all OC efferent cells were significantly smaller than those of normal hearing animals (Cantos, Lopez et al. 2000, Cantos, Lopez et al. 2003). Our previous explorations of MOC efferent cells labelled using ChAT immunohistochemistry showed that mice with prolonged deafness exhibited a progressive decrease in MOC efferent cell size as a function of length of deafness rather than age per se (Suthakar, Connelly et al. 2013). In normal hearing CBA/CaJ mice, a substantial decrease in MOC, but not LOC efferent neurons has been reported in old vs young animals (Vicencio-Jimenez, Weinberg et al. 2021). In aged gerbils, significant reductions in OC efferent counts were also observed in OC efferent populations (Radtke-Schuller, Seeler et al. 2015). We hypothesize that underlying hearing dysfunction will compound this age-related loss of OC efferent neurons.

There exists an intricate interplay between the sensory epithelia of the cochlea, and its neuronal connectivity. Indeed, knocking out the proneuronal gene (*ngn1*), which is involved in development of sensory neurons in the cochlear, results in improper embryonic axonal guidance in OC efferents, and a complete loss of innervation of hair cells (both afferent and efferent) at birth (P0) (Ma, Anderson et al. 2000). Similarly, the situation in cochlear ablation is confounded by the idea that complete destruction of mature hair cells potentially has differential down-stream effects on afferent and efferent systems. Notably, significant damage occurs to spiral ganglion cells in addition to their dendrites, whereas only the axons of OC efferents are affected. In fact, following unilateral cochlear ablation in adult rats, a strong upregulation in growth associated protein 43 (GAP-43) mRNA occurs in terminals in the ventral cochlear nucleus and cells of the VNTB, presumed to be contralateral MOC efferents (Kraus and Illing 2004). The MOC collaterals into the cochlear nucleus (Brown 1993) onto T- and D-stellate neurons (Baashar, Robertson et al. 2019) may sustain MOC plasticity capable of altering central circuitry following cochlear damage despite the loss of both their major source of afferent input and peripheral targets (Kraus and Illing 2004, Kraus, Ding et al. 2013).

The ability to understand speech in noise is one of the functions generally attributed to the OC efferent system. Because difficulties in speech understanding accompany hearing loss, we wondered whether OC efferent dysfunction might be involved and hypothesized that the extent of OC efferent dysfunction would be proportional to the severity of hearing loss. Given the traumatic nature of the cochlear ablation technique, we adopted two mutant mice models to investigate the nature and time course of OC efferent degeneration in progressive hearing loss and congenital deafness. We used retrograde tract-tracing coupled with light and fluorescent microscopic methods to investigate changes to OC efferents using quantitative anatomical methodology in mice with normal hearing, hearing loss, and congenital deafness, and expand on existing reports by probing alterations to individual cell morphology.

## Materials and Methods

### Animals

Data presented in the present study were obtained from animals used in another study published by the same authors (Suthakar and Ryugo 2017). A total of 16 adult wild type CBA/CaH, 10 DBA/2J, and 9 Shaker2-/- mice of either sex were used in this study. CBA/CaH mice were derived from Jackson Laboratory CBA/CaJ mice (strain #000654), renamed at the request of Jackson Labs, and established as an inbred colony at the Australian BioResources Facility (Mossvale, New South Wales, AUS). This strain was used as normal hearing controls as they have previously been documented to retain normal hearing until over 26 months of age (Spongr, Flood et al. 1997). The DBA/2J mouse exhibits early onset hearing loss at around 3 weeks of age due to a mutation in the cadherin-23 gene (Cdh23), which affects the tip links of stereocilia atop hair cells (Willott, Kulig et al. 1984, Hultcrantz and Spangberg 1997, Johnson, Zheng et al. 2000, Wang and Manis 2006). The homozygous shaker2 (Shaker2-/-) mouse exhibits congenital deafness due to a point mutation in the gene, Myo15, on chromosome 11, whose expression is limited to the inner ear and pituitary gland (Probst, Fridell et al. 1998, Liang, Wang et al. 1999). Functionally, the shortened stereocilia atop hair cells affect both auditory and vestibular systems producing phenotypic deafness and circling behavior (Beyer, Odeh et al. 2000, Lee, Cahill et al. 2003).

### Auditory Brainstem Responses (ABRs)

A total of 32 animals from 9 cohorts (3 strains: CBA/CaH, DBA/2J, Shaker2-/-; 3 age groups: 1, 3, and 6 months) were used for auditory brainstem response (ABR) testing. ABR testing was performed under ketamine/xylazine (50mg/kg and 10mg/kg, respectively) anesthetic in a double walled, sound attenuated chamber (Sonora Technology Co., Gotenba, Japan). Following the disappearance of the toe pinch reflex, the animal was placed on a battery-operated infrared heat pad to maintain body heat, and a lubricating ophthalmic ointment was applied to avoid drying of the eyes. Needle electrodes were inserted under the skin where the recording electrode was positioned at the vertex, the reference electrode was positioned under the right pinna, and the ground electrode was inserted into the muscle of the biceps femoris.

Free-field click (0.1ms rectangular pulse, repetition rate of 10/sec) and tone stimuli (4, 8, 16, 24, 32 and 40 kHz; 5ms duration; 0.5ms rise-fall) were generated with a signal processor (RZ6; Tucker Davis Technologies [TDT], Alachua, FL) controlled by BioSigRZ software (v5.3; TDT), preamplified (Medusa RA16PA; TDT) and delivered using a magnetic speaker (MF1; TDT) placed 10cm away from the right pinna. Stimuli were presented at 10dB decreasing steps from 90dB SPL to 0dB SPL. Averaged responses to 512 stimuli presentations were amplified (Medusa RA16PA/RA4LI; TDT), filtered (0.5-3kHz; notch at 50Hz), and recorded (RZ6; TDT) for offline analysis.

### Distortion Product Otoacoustic Emissions (DPOAEs)

The same animals described above for ABR testing were used to obtain distortion product otoacoustic emission (DPOAE) data. A miniature assembly consisting of a calibrated low noise microphone (ER 10B+, Etymotic, Chicago, IL) connected to two electrostatic speakers (EC1, TDT) via separate coupling tubes 10 cm in length (ER10B-3, Etymotic) was inserted into the right ear canal by manipulating the pinna with fine forceps.

Stimuli were presented with a signal processor (RZ6; TDT) controlled by *BioSigRZ* software (v5.3; TDT). Each electrostatic speaker presented a pure tone frequency calculated as a function of the audiometric frequency (f) being measured where the ratio of *f_2_*/ *f_1_* = 1.2, *f_1_* = 0.909 * f, and *f_2_* = 1.09 * f. Stimuli were presented from 80dB SPL to 20dB SPL in 10dB decrements. Averaged responses to 512 stimuli presentations of 20.97ms of length were used to perform a Fast Fourier Transform (FFT), resulting in a frequency-domain waveform from which the amplitude of the primary frequencies (*f_1_* and *f_2_*) and cubic distortion product (*2f_1_-f_2_*) was obtained.

### Tracer Injections

A total of 30 animals received injections of a retrograde neuronal tracer into the cochlea to label olivocochlear neurons and to study their morphology as a function of age and hearing status. Immediately after the completion of hearing tests, individual animals were transferred to an aseptic operating table. Body temperature was maintained throughout surgery using a battery-operated infrared heat pad. The animal was clamped into a custom inhalational anesthetic assembly constructed using a snout clamp (Item # 51629, Stoelting, Wood Dale, IL) and plastic tubing to deliver isoflurane (1.5-2.0% in ∼600cc/min O_2_). Additionally, light suction was employed to prevent pooling of anesthetic gas around the snout and to ensure continuous airflow. Injections of Fluorogold (FG; 4% in saline; Fluorochrome, Denver, CO) or cholera-toxin subunit-B (CTB; 0.5% in 0.5M Tris buffer, pH 7.6 and 3M NaCl; List Biological Laboratories, Campbell, CA) were made into the round window of the cochlea as previously described (Suthakar and Ryugo 2017).

### Tissue processing

The post-injection survival period depended on the dye used 9 days for FG injections and 14 days for CTB injections. At the end of this period, animals were deeply anesthetized with a lethal dose of sodium pentobarbitone and exsanguinated using a standard transcardial perfusion method using a buffered 4% paraformaldehyde solution. Heads were removed and postfixed in 4% paraformaldehyde for 2-3 hours. The brain was dissected from the skull and embedded in a mixture of gelatin (25mg) dissolved in 5ml of hot water and bovine serum albumin (1.5g) hardened with 1mL of 37% formaldehyde and 400µL of 5% glutaraldehyde. The embedded block was cut into 60µm sections in the coronal plane on a vibrating microtome (VT1200S; Leica Systems, Nussloch, FRG). All immunohistochemistry steps were performed on free-floating sections and all reagents made up in 0.12M Tris-buffered saline unless stated otherwise. Rinses were always performed thrice within a 15-min window using fresh 0.12M Tris-buffered saline.

Immunohistochemical processing steps for visualization of FG or CTB tracer were identical to those reported previously (Suthakar and Ryugo 2017). Briefly, this involved incubating sections in 1% H_2_O_2_ for 10 minutes to remove endogenous peroxidase. Sections were rinsed, incubated for one hour in 0.1% PhotoFlo (Kodak, Rochester, NY), followed by one hour in 1% normal goat serum (FG; Cat # S26-100ML; Millipore, Billerica, MA) or 1% normal rabbit serum (CTB; Cat # S-5000; Vector Laboratories, Burlingame, CA), then incubated overnight at 4°C on a shaking platform in either polyclonal rabbit anti-FG primary antibody (1:50,000; Cat# Fluorogold, RRID:AB_2314408; Fluorochrome) or polyclonal goat anti-CTB primary antibody (1:10,000; Cat# 703, RRID:AB_10013220; List Biological Laboratories). The following day, sections were washed and incubated in biotinylated goat anti-rabbit secondary (FG; 1:200; Cat# BA-1000; Vector Laboratories, Burlingame, CA) or biotinylated rabbit anti-goat secondary (CTB; 1:200; Cat# BA-5000; Vector Labs) for one hour, washed again, incubated in ABC (Vectastain Elite ABC Kit; Cat# PK-6100; Vector Labs) for another hour, and developed using nickel-intensified diaminobenzidine (Ni-DAB; FG), or diaminobenzidine (DAB; CTB). Sections were mounted and coverslipped using Permount (Fisher Scientific, Pittsburgh, PA).

Immunohistochemical processing for ChAT consisted of three rinses in 0.12M Tris buffered saline (TBS), a 10-minute incubation in 3% H_2_O_2_, and rinsed again. All sections were incubated for 1 hour in 0.1% PhotoFlo (Kodak, Rochester, NY), followed by 1 hour in 10% normal goat serum (Millipore, Billerica, MA). Sections were incubated overnight at 4°C in mouse anti-ChAT primary antibody (Vector Labs, Burlingame, CA) at a dilution of 1:1000, with 2% normal goat serum and 0.1% PhotoFlo. Negative control sections did not receive primary antibody and were instead incubated overnight in 0.12M TBS. All solutions were made in 0.12M TBS. The following day, sections were washed thrice and incubated in 1:200 biotinylated goat anti-mouse secondary antibody (Vector Labs, Burlingame, CA) for 1 hour. Sections were washed again and developed using diaminobenzidine intensified by nickel ammonium sulfate in 0.1M PBS (Tago, et al., 1989). Sections were mounted, counter stained with cresyl violet, and coverslipped for examination.

### Tissue Analysis

Sections were examined under a light microscope to verify that immunohistochemical methods stained specifically with a low background. Serial sections were examined at low magnification and those containing the SOC were gathered and blinded to the experimenter by an outside observer. Each section was photographed at low magnification, which was then used to orient sections rostro-caudally, determine the boundaries of the VNTB using a mouse brain atlas (Franklin and Paxinos, 2008), and identify labelled MOC and LOC neurons. Cells within the VNTB region (MOCs) and LSO region (LOCs) were traced using *Neurolucida* software (Microbrightfield, Essex, VT, USA) coupled to a calibrated Waccom drawing tablet. Criteria for positive identification of a cell was the observation of cellular structures such as the nucleus or dendritic projections, which were visible while focusing up and down in the Z-plane.

Measurement data were imported using Microsoft Excel (Microsoft Corporation, Redmond, WA, USA).

### Data analysis

All analyses were performed on tissue that had been blinded to the experimenters. Two different analysis strategies were utilized depending on whether the tissue was processed for brightfield or fluorescent microscopy. Brightfield tissue (CTB) was analyzed using *Neurolucida* software (MBF Bioscience, Williston, VT) attached to a light microscope (Eclipse E600, Nikon, Tokyo, Japan) controlled via a motorized stage (LEP, Ludl Electronic Products, Hawthorne, NY). Each blinded section was photographed and anatomical landmarks such as the outline of the brain, the midline, fourth ventricle, and inferior cerebellar peduncles were marked at low magnification (20x). A marker was placed on each retrogradely labelled OC efferent cell at 200x magnification, which was subsequently traced at 400-1000x magnification. To minimize photobleaching of fluorescent tissue (FG), low magnification (100-200x) photographs were used for orientation and high magnification z-stacked (400-1000x) photographs were used for offline cell counts. Eyepiece magnification was always 10x.

Statistical analyses of data were performed using *MATLAB* (v.R2013b; MathWorks, Natick, MA) and *Prism* (v.6; GraphPad Software, La Jolla, CA). Quantification of light microscopic tissue was performed using *Neurolucida* (v.11.03; MBF Bioscience). When necessary, micrographs were adjusted in *Photoshop* (v.CS4; Adobe Systems Inc., San Jose, CA) using the ‘Levels’ feature to achieve optimum brightness, contrast, and evenness of illumination across the entire image. All data values are reported as mean ± SD unless stated otherwise.

## Results

### ABRs

ABR thresholds were determined using a custom built web-based analysis program (Kirkpatrick, Ryugo et al. 2015) where threshold was defined as the sound level (dB SPL) required for detectable peak amplitude signals to be greater than 4 standard deviations above baseline noise (Bogaerts, Clements et al. 2009). Statistical analyses were performed using *Prism*. A two-way ANOVA showed significant main effects of frequency (*F*(6,58) = 33.21, p ≤ 0.0001), but not age (*F*(2,58) = 2.858, p = 0.0655) in CBA/CaH mice. In DBA/2J mice, a significant interaction (*F*(12,100) = 4.695, p ≤ 0.0001) was observed in addition to main effects of frequency (*F*(6,100) = 28.99, p ≤ 0.0001) and age (*F*(2,100) = 23.55, p < 0.0001), which was expected given the progressive age-related hearing loss phenotype in this strain. Multiple comparisons testing using Tukey’s test within frequencies, and between ages showed significant ABR threshold elevations in 6-month-old animals in response to clicks (1-month: p ≤ 0.0001; 3-month: p = 0.0004), 4kHz (3 month: p = 0.0017), 8kHz (1-month: p ≤ 0.0001; 3-month: p = 0.0263; also 1-month vs 3-month: p ≤ 0.0001), and 16kHz (1-month: p = 0.0015; also 1-month vs 3-month: p = 0.0021). No significant differences between ages were observed in DBA/2J ABR thresholds at 24-, 32- or 40-Hz. One-month-old DBA/2J mice exhibited responses comparable to CBA/CaH mice in response to Clicks and at 4 and 8 kHz, but had substantially elevated thresholds at 16, 24, 32, and 40 kHz (p ≤ 0.0001). Compared to age-matched CBA/CaH mice, 3-month-old and 6-month-old DBA/2J had significantly elevated thresholds (p ≤ 0.0001) at all frequencies except for 4 kHz. Homozygous Shaker2 mice were verified to be congenitally deaf using the Pryer’s reflex and did not exhibit neural activity to sound at any frequency at any age tested.

### DPOAEs

DPOAE data were exported as .csv files and analyses were performed offline using a combination of *Excel* (Microsoft, Redmond, WA), *MATLAB* and *Prism*. A total of 2048 data points were obtained from the FFT. For each stimulus-level parameter, the median of each set of 2048 data points was used as the noise floor (NF). DPOAE thresholds (Figure 1B) in each case were determined by plotting the rate-level curve of the magnitude (dBV) of the cubic distortion product and NF for each audiometric frequency at each level presentation. Threshold was extrapolated from plot as the level above the intersection of 2*f*_1_-*f*_2_ and NF.

**Figure 1.**
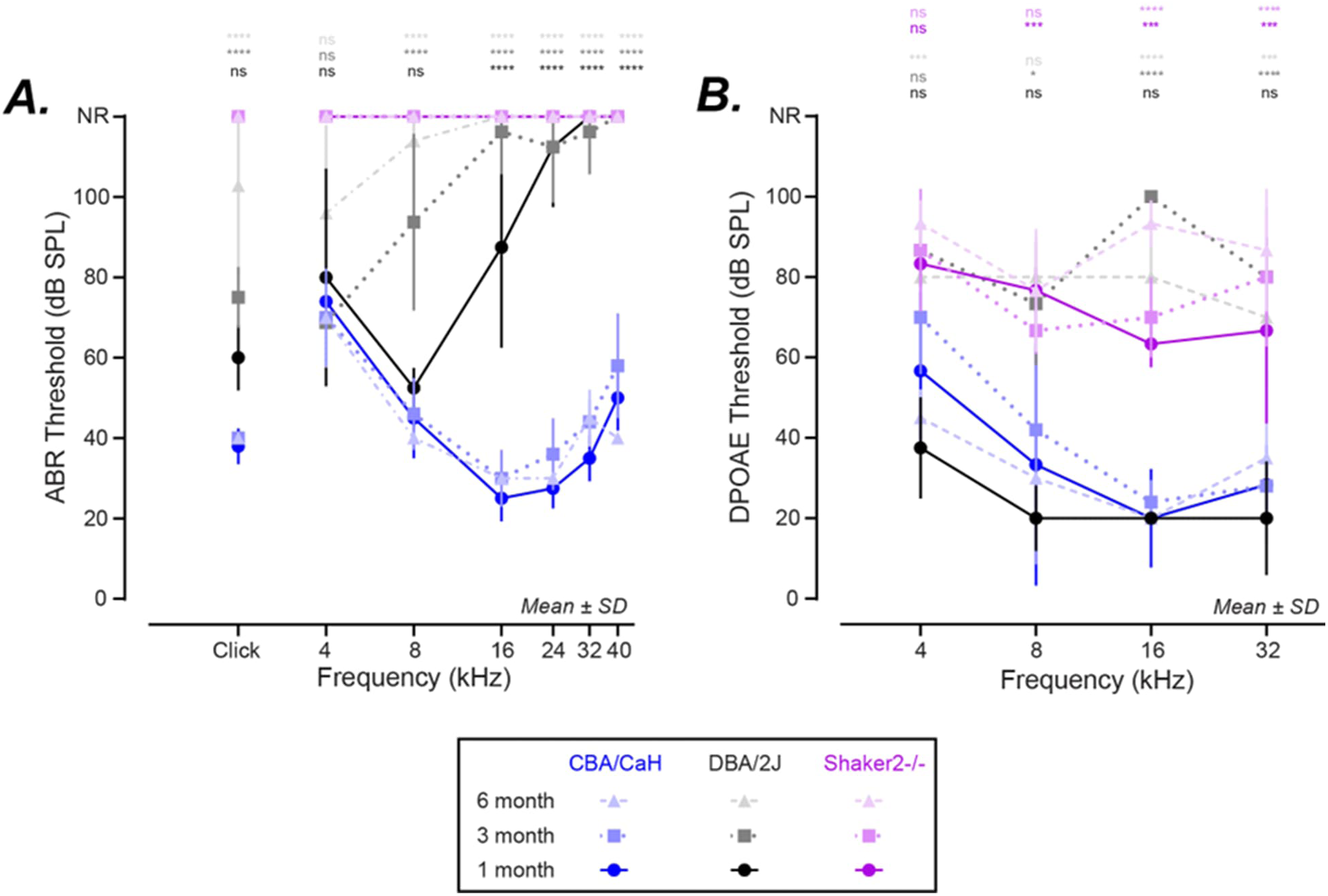
Auditory Brainstem Response (ABR) and Distortion Product Otoacoustic Emission (DPOAE) thresholds reflecting strain- and age-related cochlear function in normal hearing CBA/CaH, progressive high frequency hearing loss in DBA/2J, and congenital deafness in homozygous Shaker2-/- mice. **A.** ABR thresholds are consistent across frequencies in CBA/CaH mice at 1-, 3-, and 6-months of age. The slight increase in CBA/CaH thresholds observed at 16, 24, 32, and 40kHz were not significant (p = 0.079). DBA/2J mice exhibit elevated thresholds at all ages at 16kHz and above (p < 0.0001). The elevations in threshold progressed into the lower frequency regions with age, with thresholds significantly higher than normal at 8kHz and in response to clicks at 3- and 6-months of age (p < 0.05). The congenitally deaf Shaker2-/- mouse exhibited no neural response to sound at any age. B. No significant DPOAE threshold shifts occur as a function of age in CBA/CaH or Shaker2-/-animals. In DBA/2J mice, there was no significant change between 3- and 6-month-old animals, but both cohorts showed significantly higher thresholds than 1-month-old animals. DPOAE thresholds in 1-month-old animals showed frequency specific threshold elevations in DBA/2J (black) and Shaker2-/- (pink) strains when compared with normal hearing CBA/CaH (blue) animals. Significant threshold elevations were only observed in higher frequencies at 3 months of age. At 6 months of age, Shaker2-/- animals retained elevated thresholds across frequencies, while DBA/2J animals showed variable elevations compared to CBA/CaH mice. *\* p < 0.05; ** p < 0.01; *** p < 0.001; **** p < 0.0001*.

A two-way ANOVA showed significant effects across frequency (*F*(3,39) = 6.775, p = 0.0009) but not age (*F*(2,39) = 0.8706, p = 0.4267) in CBA/CaH mice. In DBA/2J mice, there was a significant effect of both frequency (*F*(3,25) = 3.005, p = 0.0493) and age (*F*(2,25) = 125.6, p ≤ 0.0001). A significant main effect of age (*F*(2,24) = 4.873, p = 0.0167) was observed in homozygous Shaker 2 mice. Multiple comparisons using Tukey’s test within frequencies, between ages showed significantly elevated DPOAE thresholds in 1-month-old homozygous Shaker2 mice compared to age matched DBA/2J mice (4kHz: p = 0.0023; 8kHz: p = 0.0002; 16kHz: p = 0.0072; 32kHz: p = 0.098) at all frequencies tested, and CBA/CaH mice (8kHz: p = 0.0018; 16kHz: p = 0.0026; 32kHz: p = 0.006) with the exception of 4kHz. One month old DBA/2J thresholds were not significantly different from age-matched CBA/CaH controls. Compared to 3-month-old CBA/CaH mice, DPOAE thresholds were significantly elevated at all frequencies higher than 4kHz (8kHz: p = 0.0252; 16kHz: p ≤ 0.0001; 32kHz: p = 0.0002) in DBA/2J mice, and all frequencies higher than 8kHz (16kHz: p = 0.0009; 32kHz: p = 0.0002) in homozygous Shaker2 mice. At 6 months of age, DPOAE thresholds were significantly elevated compared to age-matched CBA/CaH mice at all frequencies in both DBA/2J mice (4kHz: p = 0.0079; 8kHz: p = 0.0003; 16kHz: p ≤ 0.0001; 32kHz: p = 0.0002) and homozygous Shaker 2 mice (4kHz: p = 0.0004; 8kHz: p = 0.0006; 16kHz: p ≤ 0.0001; 32kHz: p = 0.0002).

### Distribution and qualitative morphology of labelled OC efferents

The pattern of retrograde labelling from FG and CTB injections into the cochlea was consistent across strains. FG tended to fill somata and main dendrites, whereas CTB was more clearly localized to the cell body (Figure 2). In normal hearing CBA/CaH mice, we counted on average 379.4 ± 125.7 (n = 16 animals; Table 1) labelled OC cells, comparable to previous reports in normal hearing ICR and CBA/CaJ mouse strains (Campbell and Henson 1988, Brown and Levine 2008).

**Figure 2.**
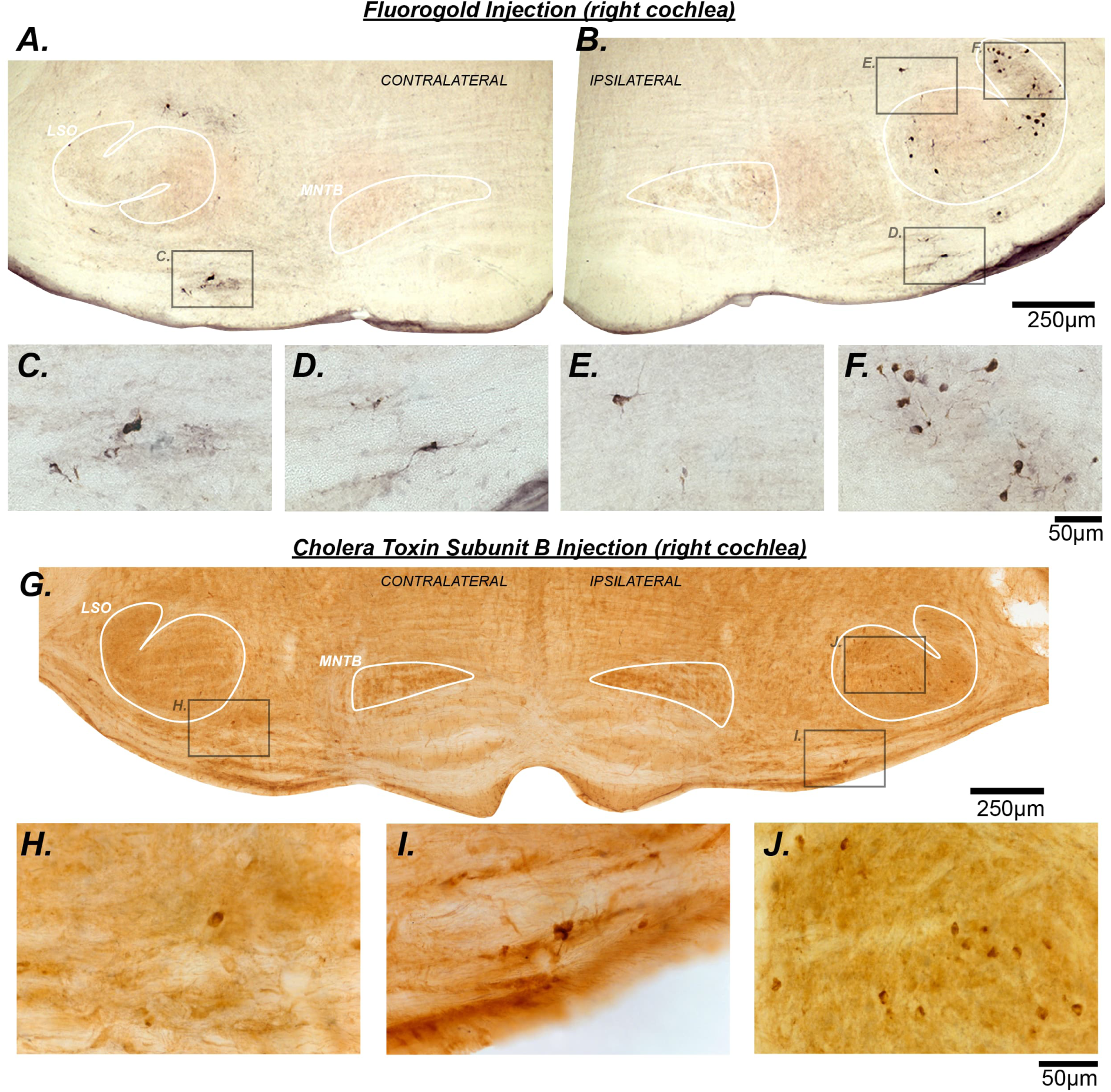
Patterns of FG & CTB staining of OC efferent neurons in a single representative coronal section of CBA/CaH mice. **A-B.** Low magnification photomicrographs of the superior olivary complex (SOC) following a unilateral injection of fluorogold tracer dye into the right cochlea. The lateral superior olive (LSO) and medial nucleus of the trapezoid body (MNTB) are indicated in grey. **C-D.** High magnification photomicrographs of MOC efferent neurons in the ventral nucleus of the trapezoid body (VNTB) indicated by insets in **A** and **B**. Contralaterally projecting MOC efferents (**C**) appear morphologically similar to their ipsilaterally projecting counterparts (**D**), both large and stellate in appearance. **E-F.** High magnification photomicrographs of a single LOC efferent ‘shell’ neuron (**E**) and small, disc shaped LOC efferent neurons located within the LSO proper (**F**). **G.** Low magnification photomicrograph of the ventral edge of the brain illustrating the superior olivary complex following a unilateral injection of choleratoxin subunit B (CTB) in the right cochlea. **H-I**. High magnification photomicrographs of MOC neurons that project contralaterally (**H**) and ipsilaterally (**I**). **J**. High magnification photomicrograph of LOC efferent neurons located within the LSO.

**Table 1.** Summary of cell counts by cohort and efferent type.

| | Age<br>(months) | n | LOC<br>(mean ± SD) | MOC<br>(mean ± SD) | Total<br>(mean ± SD) |
| --- | --- | --- | --- | --- | --- |
| **CBA/CaH** | 1 | 6 | 216.5 ± 73.7 | 145 ± 50.8 | 370.2 ± 90.0 |
| | 3 | 5 | 251.6 ± 97.1 | 219.8 ± 60.4 | 483.8 ± 129.7 |
| | 6 | 5 | 138.0 ± 56.1 | 140.0 ± 53.2 | 286.0 ± 107.9 |
| **DBA/2J** | 1 | 2 | 175.5 ± 116.7 | 166.0 ± 103.2 | 342.5 ± 13.4 |
| | 3 | 3 | 250.7 ± 71.2 | 187.7 ± 104.2 | 442.7 ± 180.4 |
| | 6 | 2 | 169.5 ± 14.8 | 113.0 ± 1.4 | 285.0 ± 11.3 |
| **Shaker2-/-** | 1 | 2 | 243.5 ± 10.6 | 77.5 ± 5.0 | 326.0 ± 8.5 |
| | 3 | 2 | 116.0 ± 22.6 | 49.5 ± 2.1 | 166.5 ± 21.9 |
| | 6 | 3 | 77.7 ± 31.1 | 24.7 ± 20.6 | 104.0 ± 11.4 |

The VNTB and the adjacent rostral periolivary complex (RPO), on both sides contained large, stellate cells with extensive dendritic projections morphologically identified as MOCs (Figure 2 A-D, G-I; Figure 3). Small disc-shaped cells were labelled within the lateral superior olive and identified as ‘intrinsic’ LOCs efferents (Figure 2 A-B, F-G, J; Figure 3). Proximal dendrites were poorly labelled in fluorescent tissue, although the somata of these cells have a distinct spindle-like appearance when viewed in the coronal plane with an orientation that seems to parallel the trajectory of incoming as well as outgoing axonal projections (Willard and Ryugo 1983, Doucet and Ryugo 2003). ‘Shell’ LOCs were also identified (Figure 2 B & E), situated around the periphery of the LSO, but because they comprise less than 5% of the total LOC population, we have combined these two populations for analysis. A previously described population of larger, stellate cells were also occasionally observed in the dorsal periolivary regions, but were not included for analysis here (DPO; see Fig. 2 of Brown and Levine 2008).

**Figure 3.**
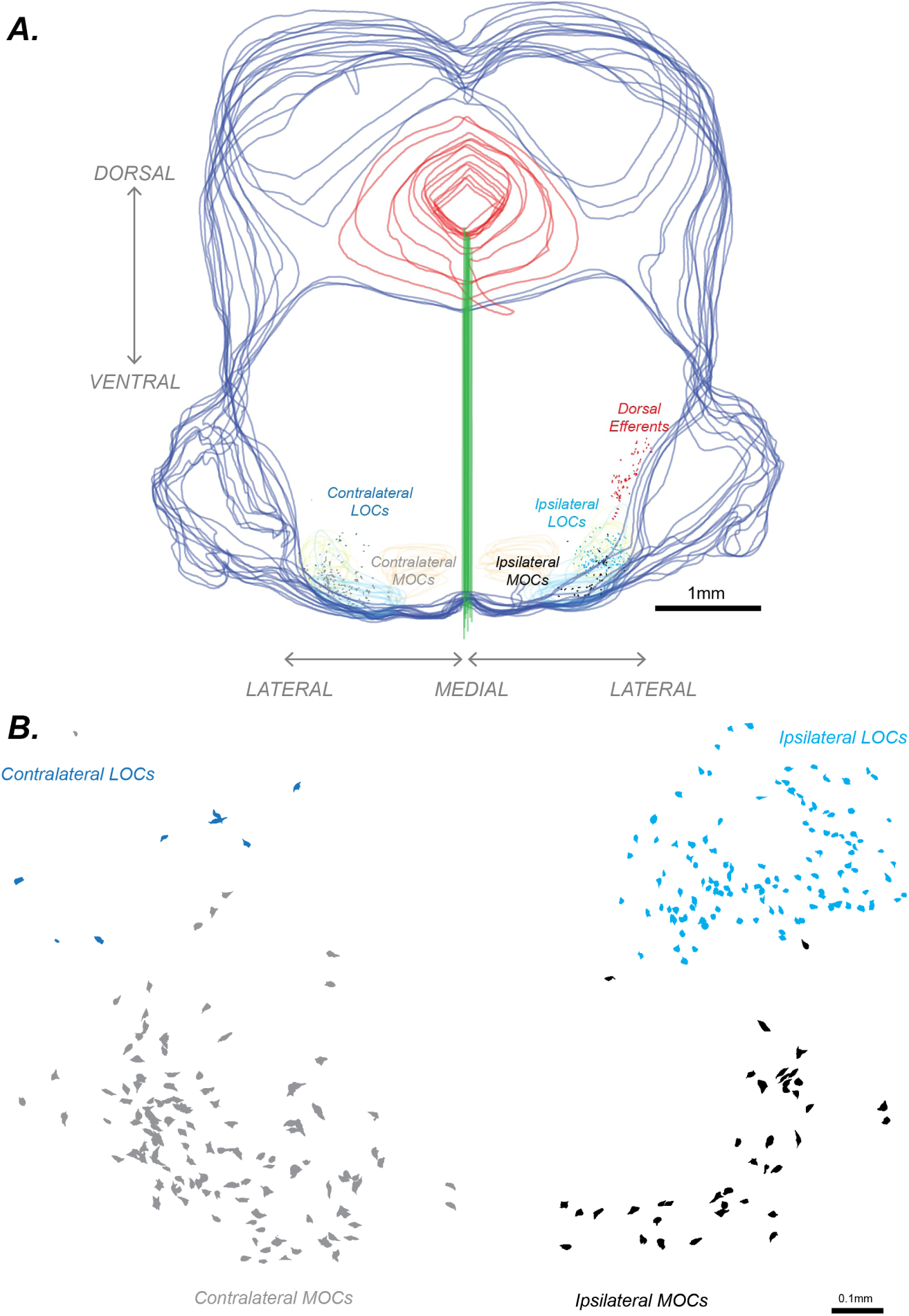
Neurolucida tracings of the brain and individual neurons from a CTB injected CBA/CaH mouse. **A.** Example of traced sections and neurons from a single 3-month-old CBA/CaH case which have been overlaid to visualize the anterior-posterior view looking through the brain. The rostrocaudal (z) axis has been flattened. The entire brain is outlined in blue, and the fourth ventricle is outlined in red. The vertical green line from the floor of the fourth ventricle identifies the midline of the brainstem. The LSO are outlined in lime green, the VNTB in light blue and MNTB in orange. Each traced cell is color coded by identity (LOC or MOC) and projection pattern (Contralateral or Ipsilateral). Dorsal efferent neurons that we traced but not included in this study are shown in red for reference. **B.** Enlargements of traced cells demonstrate qualitative differences in morphology and distributions between contralateral MOCs (grey fill), contralateral LOCs (dark blue fill), ipsilateral MOCs (black fill), ipsilateral LOCs (light blue fill) across the mediolateral axis.

**Figure 4.**
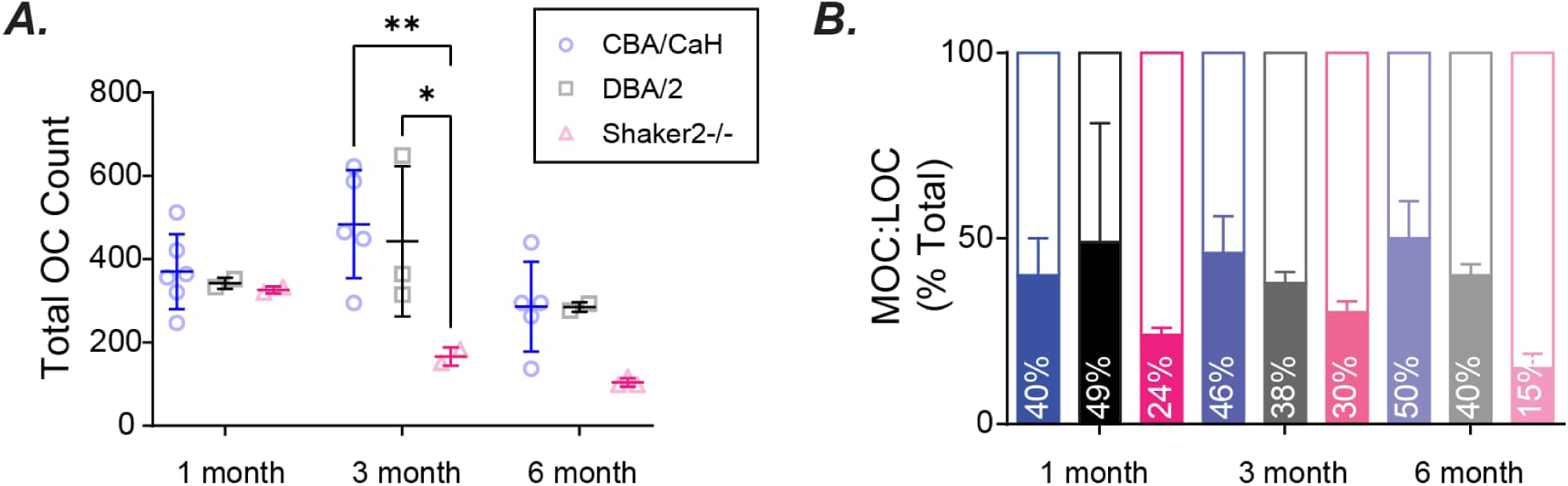
Total olivocochlear (OC) neuron counts and proportions. **A.** Total OC efferent neuron counts grouped by age and strain. No significant difference in cell counts were observed across strains at one month of age, indicating that OC efferents appear to develop normally irrespective of hearing status. By three months of age, Shaker2-/- mice had significantly fewer cells compared with age matched CBA/CaH and DBA/2J animals. **B.** The ratio of MOC to LOC efferent cells across strains were consistent with previous reports. At each age, Shaker2-/- mice had greater proportions of LOC neurons, compared with MOC neurons.

### OC efferent cell counts and distributions

Raw cell counts between staining methods were consistent and pooled for statistical analysis (Figure 3). A two-way ANOVA showed that total cell counts varied significantly across both strains (*F*(2,21) = 7.636, p = 0.0032) and age (*F*(2,21) = 4.702, p = 0.0205) (Figure 3A). Tukey’s multiple comparisons test indicated that between strain variations were only observed in the 3-month-old cohorts, showing significantly fewer OC neurons in Shaker2-/- mice compared to both CBA/CaH (p ≤ 0.01) and DBA/2J (p ≤ 0.05) animals. Importantly, no significant difference was observed between strains at one month of age, indicating that central development of OC efferent neurons appears normal.

To determine if the change in total cell count observed in the 3-month-old Shaker2-/- was attributable to either the MOC or LOC efferent neuron population, the proportion of MOC neurons was calculated across all cohorts by dividing the MOC count by the total OC count per animal (Figure 3B). A significant main effect of strain (*F*(2,21) = 5.084, p = 0.0158) was observed, with Tukey’s multiple comparisons test showing significantly lower proportions of MOCs in Shaker2-/- mice compared with CBA/CaH mice across ages (p = 0.0122). Total cell counts were divided into their respective LOC (Figure 5) and MOC (Figure 6) populations to further examine the source of the cellular loss observed in Shaker2-/- animals.

**Figure 5.**
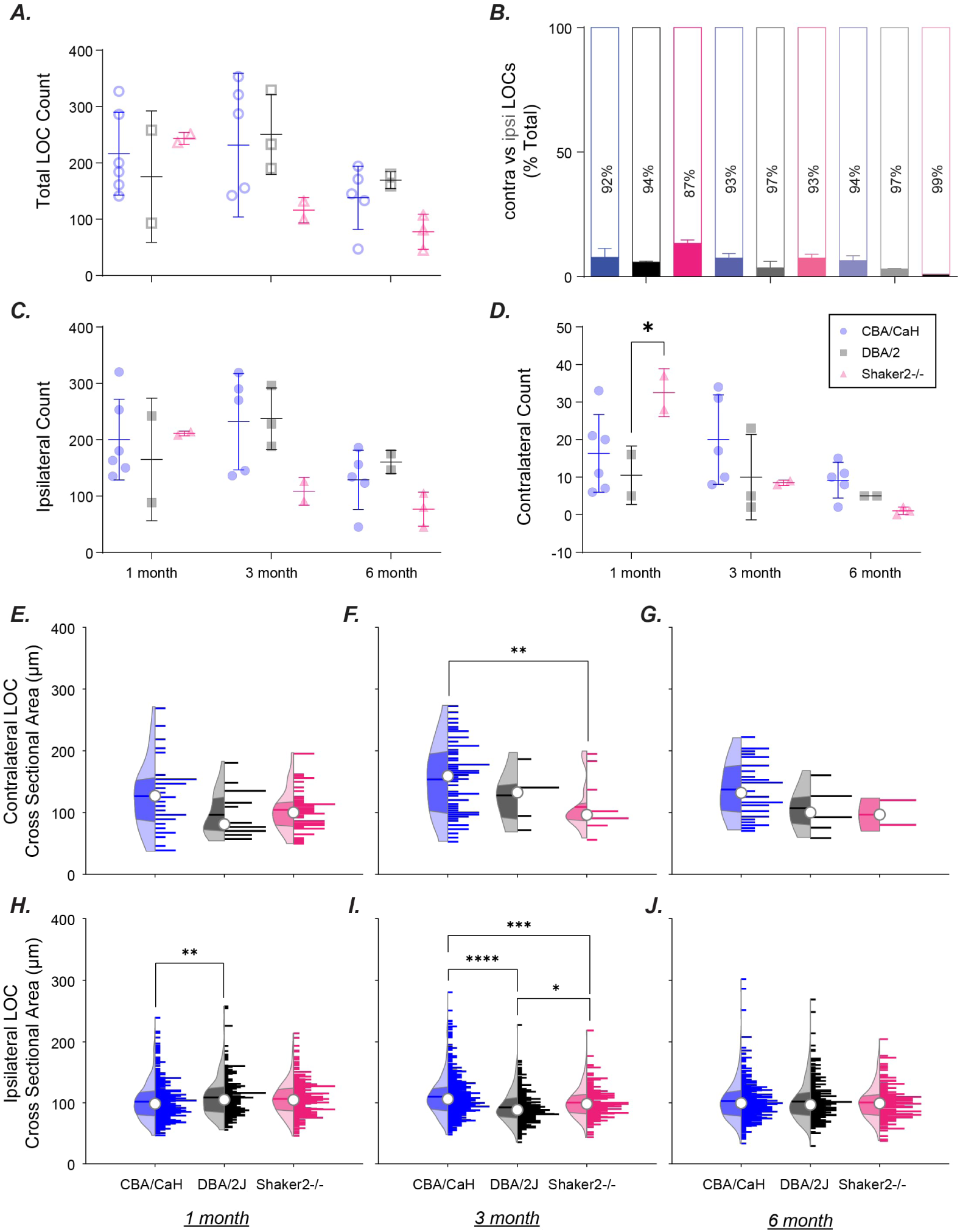
Lateral Olivocochlear (LOC) efferent cell counts, proportions and somatic cross-sectional area across strains and ages. **A.** Despite trending towards lower LOC counts with increasing age, there was no significant change in number of LOC efferents as a function of either strain or age. **B.** The proportions of contralaterally to ipsilaterally projecting LOC efferents were comparable in CBA/CaH or DBA/2J mice across ages. One-month-old Shaker2-/- mice had a greater proportion of contralaterally projecting LOCs, with numbers drastically reducing by 3-and 6-months of age. **C.** Decreases in ipsilaterally projecting LOC efferents were not statistically significant. **D.** Greater numbers of contralaterally projecting LOC efferents in one month old Shaker2-/- were observed. *Open symbols represent ‘total’ cell counts, and filled symbols represent either contralateral or ipsilateral subgroups of MOCs or LOCs.* **E-G.** Cross sectional area of contralateral LOCs at 1- (**E**), 3- (**F**), and 6- (**G**) months of age. Cross sectional area of contralateral LOCs at 1-month was variable across strains, but not statistically significant. At 3-months of age, contralateral LOCs in Shaker2-/- mice were significantly smaller than normal hearing counterparts. No significant differences in size were observed between strains at 6 months of age. **H-J.** Cross sectional area of ipsilateral LOCs at 1- (**J**), 3- (**K**), and 6- (**L**) months of age. One month old DBA/2J mice had significantly larger ipsilateral LOCs than age matched CBA/CaH and Shaker2-/- mice. At 3-months, CBA/CaH ipsilateral LOC neurons were significantly larger than age matched DBA/2J and Shaker2-/- animals. No significant differences in ipsilateral LOC cross sectional area were observed at 6-months. *The distribution on the left side of each violin plot reflects the kernel density distribution, open circle is median, horizontal line is mean, dark shading is interquartile range and light shading is range. The right side is the histogram of underlying data. * p < 0.05; ** p < 0.01; *** p < 0.001; **** p < 0.0001*.

**Figure 6.**
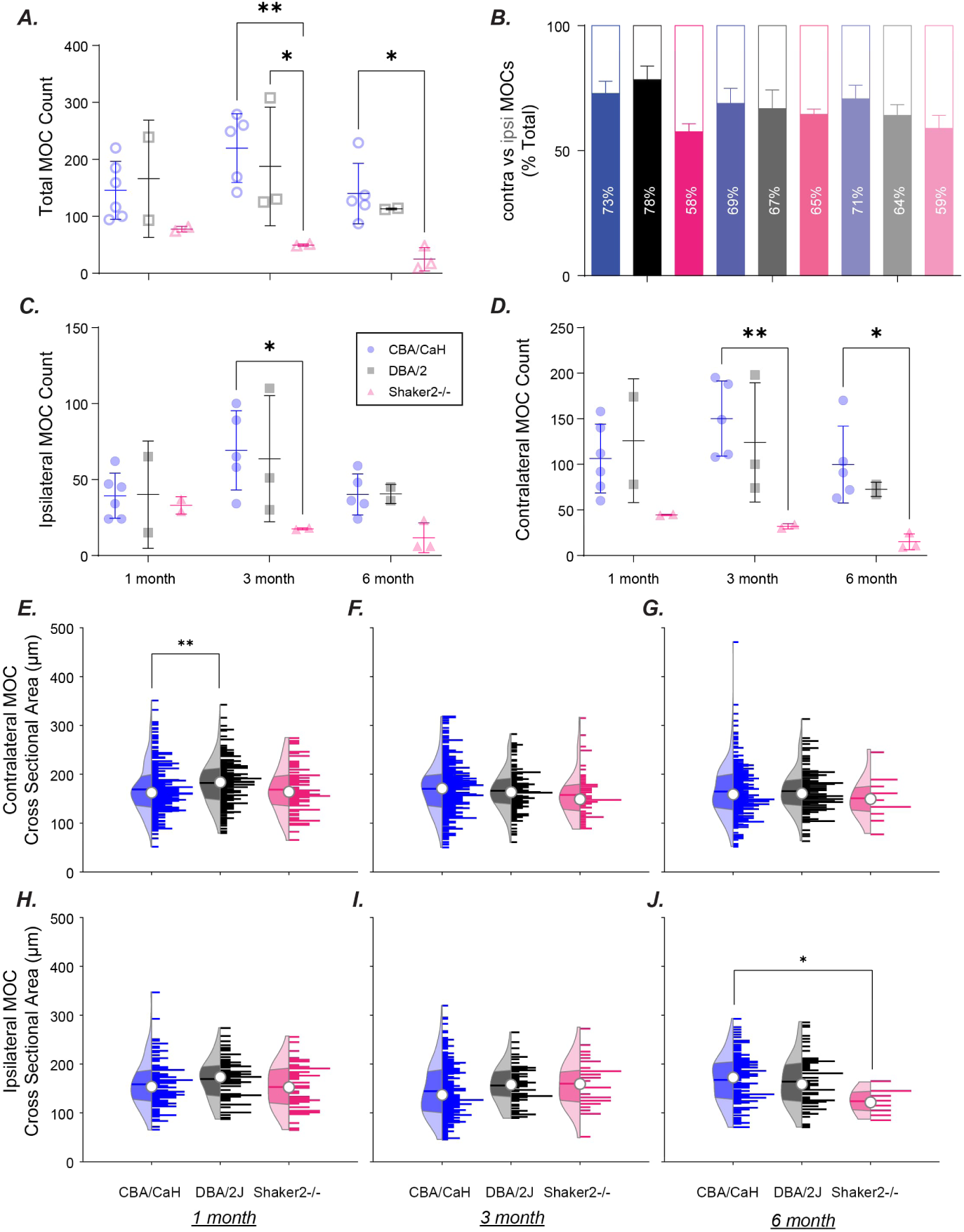
Medial Olivocochlear (MOC) efferent cell counts, proportions and somatic cross-sectional area across strains and ages. **A.** The total numbers of MOC efferents observed in Shaker2-/- mice was lower compared to both CBA/CaH and DBA/2J animals at all ages, which was statistically significant in 3-month-old animals, and 6-month-old animals (compared to CBA/CaH only). **B.** The distribution of contralaterally vs ipsilaterally projecting MOCs in normal hearing CBA/CaH and DBA/2J mice with high frequency hearing loss matched previous reports. In congenitally deaf Shaker2-/- animals, the proportion of contralaterally projecting MOCs was consistently smaller. **C.** Shaker2-/- mice had fewer ipsilaterally projecting MOCs than both CBA/CaH and DBA/2J animals. **D.** Contralateral MOC counts were comparable between CBA/CaH and DBA/2J animals at all ages, while Shaker2-/- mice had significantly fewer contralaterally projecting MOC efferents than CBA/CaH at 3- and 6-months of age. *Open symbols represent ‘total’ cell counts, and filled symbols represent either contralateral or ipsilateral subgroups of MOCs or LOCs.* **E-G.** Distributions of contralateral MOC cross sectional area are comparable between strains at 1- (**E**), 3- (**F**) and 6- (**G**) months of age. The mean cross-sectional area of 1-month DBA/2J contralateral MOCs was significantly larger than age matched CBA/CaH controls. Additionally, within the DBA/2J cohort, the 1-month-old contralateral MOCs were significantly larger than their 3- or 6-month-old counterparts. Despite large reductions in cell counts in 3-month-old Shaker2-/- mice, no differences in cross-sectional area of contralateral MOCs were observed between strains at 3-months. The drastic loss of contralateral MOC neurons in 6-month-old Shaker2-/- mice was not reflected in changes to cross sectional area. **H-J.** Cross sectional area of ipsilateral MOC neurons at 1- (**H**), 3- (**I**), and 6- (**J**) months of age. Cross sectional area of ipsilateral MOC neurons at 1-month of age are comparable across strains. Despite the significant reduction in numbers of ipsilateral MOCs in Shaker2-/-, no significant reduction in cross sectional area was observed. Unlike the situation observed in contralateral MOCs, ipsilateral MOCs in 6-month old Shaker2-/- are significantly smaller than their normal hearing CBA/CaH counterparts. *The distribution on the left side of each violin plot reflects the kernel density distribution, open circle is median, horizontal line is mean, dark shading is interquartile range and light shading is range. The right side is the histogram of underlying data. * p < 0.05; ** p < 0.01; *** p < 0.001; **** p < 0.0001*.

### Quantification of changes to LOC efferent neurons

#### LOC cell counts and distributions

A two-way ANOVA showed that LOC cell counts varied with age (*F*(2,21) = 3.730, p = 0.0411) but not strain (*F*(2,21) = 1.610, p = 0.2235) (Figure 5A). Again, in agreement with previous reports, a predominant ipsilateral projection pattern in the LOC system was observed with the ratio of contralateral to ipsilaterally projecting LOCs around 7%:93% (Figure 5B). Ipsilateral LOC cell counts (Figure 5C) remained stable over time in both DBA/2J and CBA/CaH mice. Interestingly, at 1 month of age, greater numbers of contralaterally projecting LOC neurons (Figure 5D) were observed in Shaker2-/- mice, which was significant compared to age-matched DBA/2J (p ≤ 0.05) mice, but not CBA/CaH mice.

#### Cross sectional area in LOC efferent neurons

Cross sectional area of contralateral LOCs (Figure 5E-G) varied by strain (F(2,221) = 6.774; p = 0.0014), but not age (F(2,221) = 2.526; p = 0.0823). Across all ages, contralateral LOCs in CBA/CaH mice were larger than DBA/2J mice (p = 0.0165) and Shaker2-/- mice (p = 0.0162). Cross sectional area of ipsilateral LOCs (Figure 5H-J) varied with both strain (F(2,3153) = 4.471; p = 0.0115), and age (F(2,3153) = 6.640; p = 0.0013). Within strains, significant effects were observed at all ages in DBA/2J mice; ipsilateral LOCs were significantly larger in 1-month-old (109.0 ± 32.18) compared to both 3-month-old (92.6 ± 25.41; p < 0.0001) and 6-month-old (102.0 ± 32.89; p = 0.0006) DBA/2J cohorts. Ipsilateral LOCs in 3-month-old DBA/2J mice were also smaller than those observed in their 6-month-old counterparts (p = 0.0227). Within the CBA/CaH strain, ipsilateral LOCs were larger in 3-month-old animals (110.0 ± 37.77) compared to both 1-month-old (101.9 ± 30.08; p < 0.0001), and 6-month-old animals (102.9 ± 32.96; p = 0.0004). No change in ipsilateral LOC size was observed in Shaker2-/- animals at any age.

### Quantification of changes to MOC efferent neurons

#### MOC efferent cell counts and distributions

There were significantly fewer MOC efferent neurons in Shaker2-/- mice compared to CBA/CaH controls (p ≤ 0.01) and DBA/2J mice (p ≤ 0.05) at 3-months of age (Figure 6A). At 6 months Shaker2-/- mice had fewer MOC neurons than CBA/CaH controls (p ≤ 0.01) but not DBA/2J mice with progressive high frequency hearing loss (Figure 3A). The ratio of contralateral-to-ipsilateral MOC projecting neurons was consistent with previous descriptions. In agreement with previous reports (Campbell and Henson 1988, Brown and Levine 2008), there was a greater contralateral projection in the MOC system, the ratio in normal hearing animals was around 70%:30% (Figure 6B). In normal hearing CBA/CaH animals at all ages, the contralateral-to-ipsilateral ratios (Figure 3D) matched the ranges previously described in mouse using FG (0.60), horseradish peroxidase (HRP; 0.53), and AChE staining (0.44; calculated from Brown and Levine 2008). At one month of age the contra:ipsi ratio of MOCs in Shaker2-/- mice was significantly skewed towards the ipsilaterally projecting MOCs compared to age- matched CBA/CaH (p ≤ 0.05) and DBA/2J (p ≤ 0.01) animals. No such difference was observed between strains at any other age; however, there was a general trend observed in DBA/2J animals of decreasing proportions of contralaterally projecting MOCs with advancing age/elevated thresholds. Ipsilaterally projecting MOC counts (Figure 6C) were lower in Shaker2-/- mice compared to CBA/CaH controls (p ≤ 0.05) at 3-months of age. Shaker2-/- mice had significantly fewer contralaterally projecting MOCs (Figure 6D) than CBA/CaH controls at 3-months (p ≤ 0.01) and 6-months (p ≤ 0.05) of age.

#### Cross sectional area of MOC efferent neurons

Given the drastic changes to the overall count of cells in mice with progressive hearing loss and congenital deafness, we next investigated changes to OC efferent morphology. Cross-sectional area was used to infer neuronal health vis-a-vis hearing ability and age (Figure 6E-J). The cross-sectional area of contralateral MOC neurons (Figure 6E-G) varied as a function of age (*F*(2,1626) = 4.5; p = 0.0113) but not strain (*F*(2,1626) = 2.42; p = 0.0892). Multiple comparison testing using Tukey’s test within strains and between ages showed that this effect was driven by changes observed in DBA/2J mice. Significantly larger contralateral MOC neurons were observed in the 1-month-old (mean ± SD: 182.3 ± 46.03) compared to the 3-month-old (mean ± SD: 166.1 ± 41.12; p = 0.0064) and 6-month-old DBA/2J cohorts (mean ± SD: 165.6 ± 43.77; p = 0.0075). Contralateral MOCs from the 1-month DBA/2J cohort were also significantly larger than age matched CBA/CaH mice (p = 0.0099). As expected, no significant age effects were observed in CBA/CaH mice (1-month: 169.1 ± 50.14; 3-month: 170.1 ± 50.01; 6-month: 164.5 ± 53.27). Unexpectedly, despite the drastic loss of neurons observed, no significant changes were observed in cross sectional area of contralateral MOCs in Shaker2-/- mice (1 month: 168.5 ± 47.75; 3-month: 157.8 ± 41.12; 6-month: 150.7 ± 43.61).

A small, but significant effect of strain (*F*(2,823) = 3.092; p = 0.0460), but not age (*F*(2,823) = 1.289; p = 0.2761) was observed in cross-sectional area of ipsilateral MOCs (Figure 6H-J). Within strains, 3-month-old CBA/CaH mice (144.5 ± 54.14) had significantly smaller ipsilateral MOCs than their 1-month-old (158.5 ± 48.76; p = 0.0398) and 6-month-old (167.5 ± 49.04; p < 0.0001) counterparts. No significant effects were observed in DBA/2J (1 month: 169.4 ± 41.55; 3-month: 156.2 ± 38.87; 6-month: 164.0 ± 49.82) or Shaker2-/- (1 month: 153.0 ± 45.15; 3-month: 159.8 ± 47.68; 6-month: 123.9 ± 25.4) cohorts. At 6-months of age, ipsilateral MOCs of Shaker2-/- were significantly smaller than age matched CBA/CaH animals (p = 0.0277), but it should be noted that only 9 ipsilateral MOCs were observed across 3 animals.

#### MOC efferent cell locations

Existing evidence suggests that the central locations of MOC efferents in the VNTB are tonotopically organized with respect to the frequency of their projections into the cochlea (Guinan, Warr et al. 1984, Robertson, Anderson et al. 1987, Brown 1993). Descending projections to MOC neurons from the IC are tonotopically organized with higher frequency projections located laterally, and lower frequency projections located more medially (Caicedo and Herbert 1993, Malmierca, Le Beau et al. 1996, Suthakar and Ryugo 2017). Previous data also suggests that this spatial mapping of descending frequency information is more disorganized in congenitally deaf Shaker 2-/- mice than DBA/2J mice that retained hearing ability at lower frequencies (Suthakar and Ryugo 2017).

To determine if the stark MOC efferent degeneration observed here occurred in a location specific manner, three dimensional coordinates of each MOC efferent neuron were first normalized in the mediolateral axes to compare between cases (Figure 7). For each case, correction factors were calculated using conspicuous landmarks visible in each section – the midline and inferior cerebellar peduncle (ICP; Figure 7A) which shifted slightly in rostral vs caudal sections (Figure B). X-correction factors were calculated by computing the orthogonal distance between the ICP and midline. The x-coordinate of each MOC neuron was then divided by its section x-correction factor and then multiplied by the global x-correction factor across all cases (Figure 7C). MOC location data were subsequently plotted as a function of normalized mediolateral distance (corrected and normalized x-coordinate) and rostrocaudal distance (based on cut section thickness) for each cohort (Figure 8). For each age group, MOCs were separated by ipsilateral (grey markers) or contralateral (colored markers) projection.

**Figure 7.**
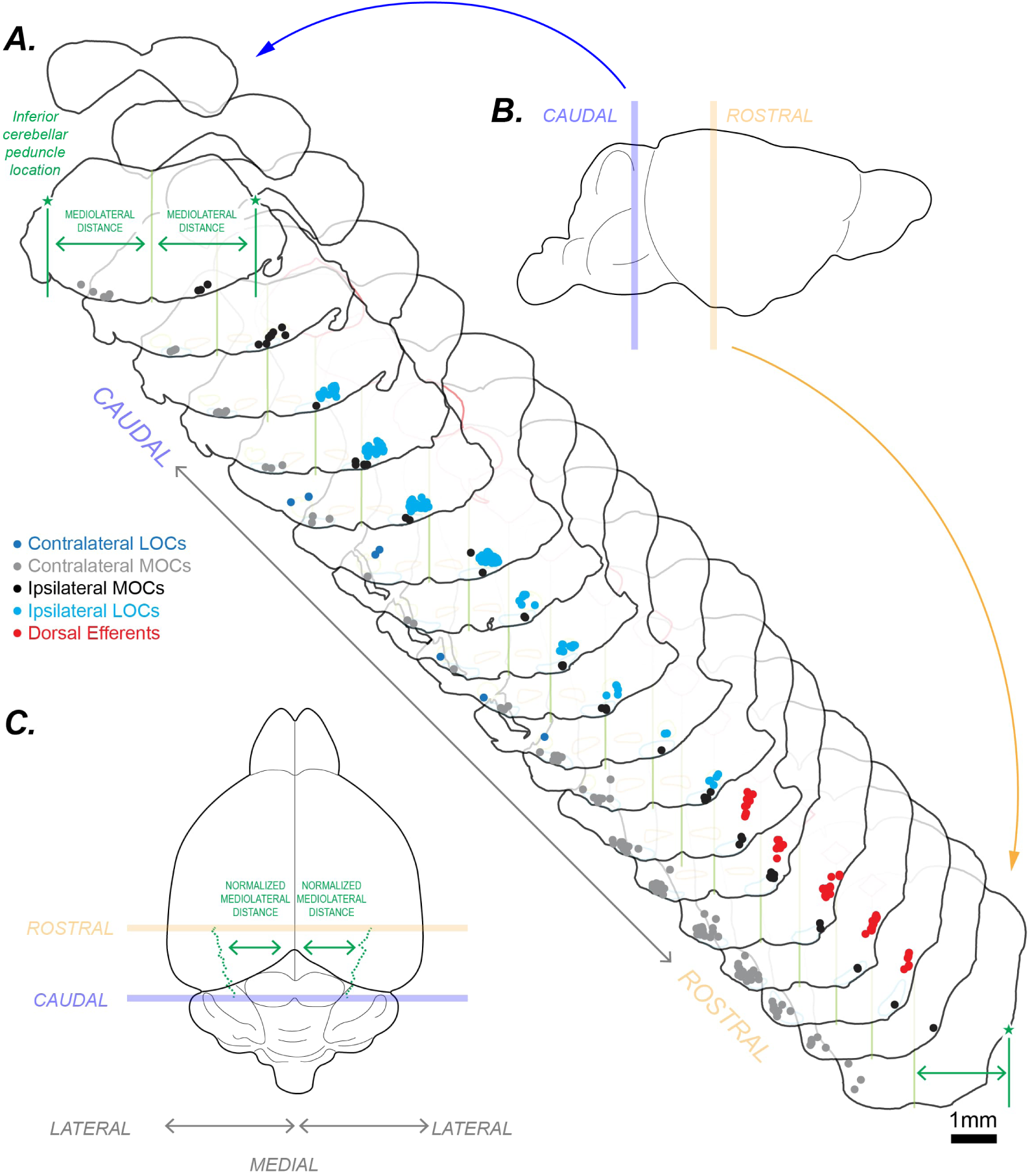
Neurolucida tracings illustrate the anatomical locations and normalization procedure for mapping MOC efferent cell locations. **A.** The same traced sections from Figure 3 have been arranged to illustrate the shifting morphology of the brain from rostral to caudal sections. MOC neurons shift from a ventromedial location in caudal sections to more dorsolateral locations in rostral sections. **B.** The approximate locations of the caudal and rostral sections have been indicated on the lateral aspect of the brain. **C.** The rostrocaudal and normalized mediolateral axes are illustrated on this superior view of the brain. To enable comparisons of MOC locations across different cases, the mediolateral location (x-coordinate) of each traced cell was normalized using consistent and easily identifiable landmarks – the midline and the inferior cerebellar peduncles. The light green vertical line identifies the midline of the brainstem, and the dotted dark green line represents the shifting location of the inferior cerebellar peduncle in different sections.

**Figure 8.**
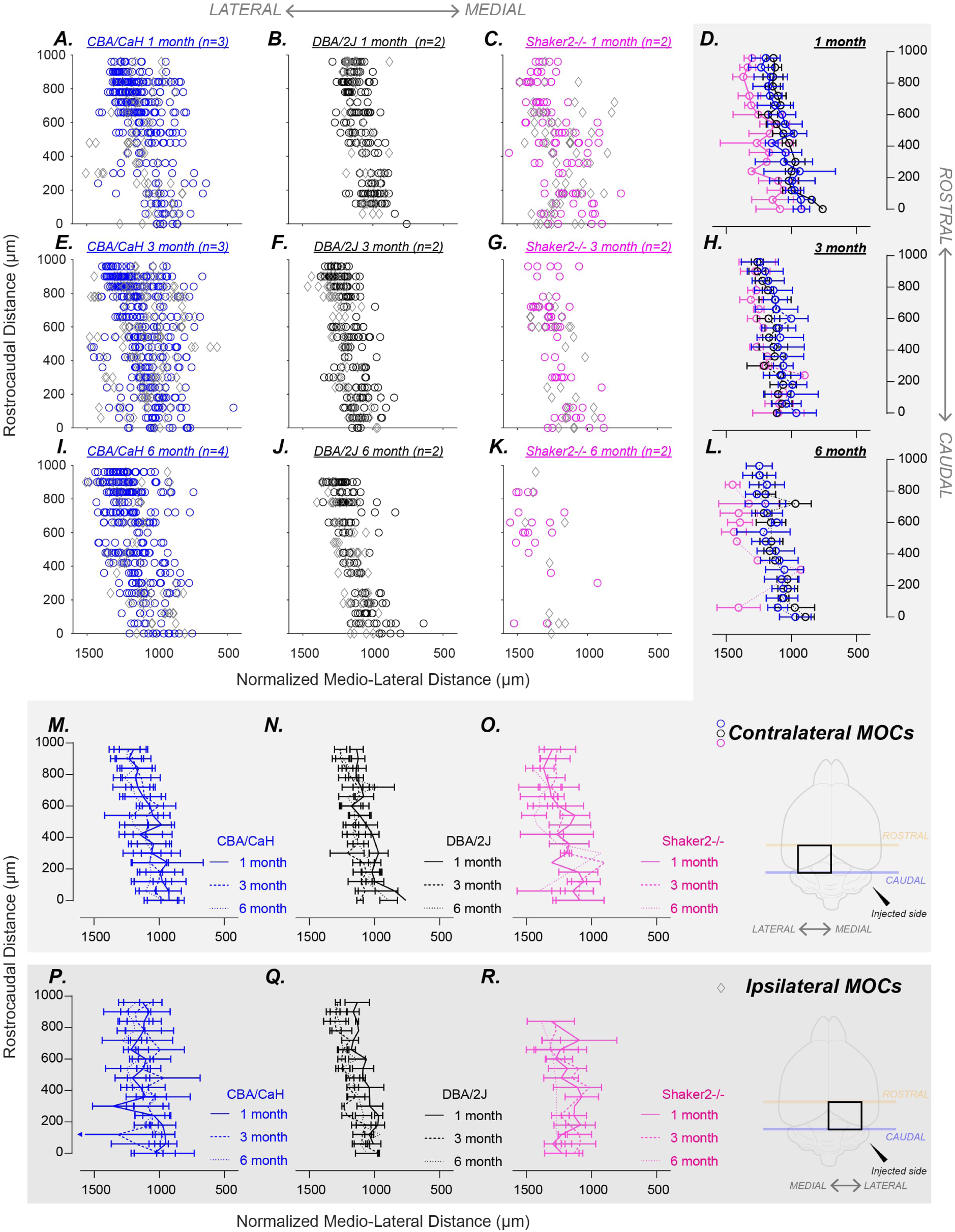
Comparison of normalized MOC efferent locations across the rostrocaudal and mediolateral axes. **A-C, E-G, I-K.** Individual MOC efferent cell locations in the VNTB were normalized along the mediolateral axis using anatomical landmarks to allow for comparison between cases. In each panel, MOCs are color coded to represent ipsilateral (grey diamond) and contralateral (colored circles) projecting MOCs. Each column refers to a strain: CBA/CaH (blue; A,E,I), DBA/2J (black; B,F,J), Shaker2-/- (pink; C,G,K), and each row refers to an age (1-, 3- or 6-month old). Within strains and across ages, the locations of ipsilateral and contralaterally projecting MOCs are comparable. In each cohort the rostrolateral and mediolateral distribution of ipsilateral and contralaterally projecting MOCs are comparable. The loss of both contralateral and ipsilateral MOCs in 6-month old Shaker2-/- mice is striking. **D,H,L & M,N,O.** Mean ± S.D of normalized mediolateral distance of contralateral MOCs for each rostrocaudal section grouped by age: 1-month (D), 3-month (H), and 6-month (L), or strain: CBA/CaH (M), DBA/2J (N), and Shaker2-/- (O). An inset box (black) overlaid on the top-down brain schematic brain illustrates the approximate location of the plots with respect to the contralateral injection site (black triangle). **P,Q,R.** Mean ± S.D. of normalized mediolateral distance of ipsilateral MOCs grouped by strain: CBA/CaH (P), DBA/2J (Q), and Shaker 2-/- (R). The ipsilateral plots have been ‘flipped’ or ‘reflected’ at the midline and presented as mirror images to enable comparison with contralaterally projecting neurons.

Qualitatively, compared to the normal hearing CBA/CaH cohort, both contralateral and ipsilateral MOCs in DBA/2J animals were confined to a narrower region in the mediolateral axis across ages (Figure 8 A-B, E-F, I-J). Cell distributions in Shaker2-/- were comparable to age-matched CBA/CaH at 1 month and began to resemble DBA/2J distributions (i.e. become narrower) at 3 months (Figure 8 C, G). By 6 months of age, the significant loss of both contralateral and ipsilateral MOCs is apparent (Figure 8 K).. To quantify these changes, we performed a one-way ANOVA between cohorts using the variance in the normalized mediolateral spread of MOCs (i.e. standard deviations) of each rostrocaudal section was performed. Variance was significantly different across age-matched cohorts for both contralateral MOCs (F(4.016,70.79) = 6.336; p = 0.0002) and ipsilateral MOCs (F(2.927,44.27) = 8.961; p = 0.0001). Follow-up multiple comparisons testing using Sidak’s test showed that the standard deviations of the contralateral MOC locations were significantly smaller in DBA/2J cohorts at 1-month (p = 0.0105), 3-month (p = 0.0072) and 6-months (p = 0.0004) of age compared to their age matched normal hearing CBA/CaH counterparts. Similarly, for ipsilateral MOCs, DBA/2J cohorts had significantly smaller standard deviations compared to age- matched CBA/CaH cohorts at 1-month (p = 0.0004), 3-month (p = 0.0325) and 6-months (p = 0.0013) of age. In congenitally deaf Shaker2-/- animals, the variance was significant for contralateral MOCs from the 6-month-old (p = 0.037) CBA/CaH cohort only, likely due to the sheer number of MOC neurons lost. No significant differences in standard deviations of normalized medio-lateral distance were observed between age matched DBA/2J or Shaker 2-/- cohorts.

## Discussion

We examined the nature of changes to OC efferent neurons in the mouse auditory brainstem based on age and severity of hearing pathology. OC efferent neurons constitute the functional effectors of the descending central auditory pathway. When the OC efferent pathway is compromised, feedback from the brain can no longer be provided to the periphery, and the descending system must rely on central feedback loops for modulation of ascending acoustic information. Therefore, changes to OC efferent pathways with respect to hearing ability and age are important considerations in assessing central mechanisms of hearing pathology. We are especially aware that efferents subserve functions beyond exacting pure tone threshold shifts in noise. Here, we have shown that a decrease in the number of OC efferents accompanies long-term congenital deafness. Secondary degeneration caused by hearing loss appears to affect the separate populations of OC efferents in a differential manner. Most notably, congenital deafness results in a selective loss of contralaterally projecting MOC efferents, and these cells exhibit a selective size reduction as reflected by cross-sectional area. In contrast, long-term acquired (and progressive) hearing loss did not appear to systematically affect the size or number of OC efferent neurons. The location of MOC neurons in the VNTB was largely comparable across strains, ages and projection patterns (i.e. contralateral vs ipsilateral). The significant loss of MOC neurons in aged congenitally deaf mice likely resulted in the marked disorganization of descending inputs from the inferior colliculus previously reported (Suthakar and Ryugo 2017).

Cell counts and size quantifications are reliable parameters to quantify neuronal changes. From an evolutionary perspective, higher cell counts are linked to increased processing complexity (Haug 1987, Flood and Coleman 1988). In fact, comparisons of auditory efferent neuron counts across taxonomic classes support this notion, with greatest efferent density observed in mammals, followed by birds, then reptiles, amphibians, and lastly, fish (Roberts and Meredith 1992). Cell size is a quantitative measure often utilized in anatomical investigations to characterize cell groups, to distinguish between healthy and atrophic neuronal populations, and to describe the degree of neuronal degeneration (e.g. Conlee, Parks et al. 1984, Born and Rubel 1985). Neuronal losses are often observed in cognitive disorders such as Alzheimer’s disease and schizophrenia (Finch 2003, Kreczmanski, Heinsen et al. 2007).

Morphological changes have been documented both peripherally and centrally with age-related hearing loss. Peripherally, degeneration has been observed in the stria vascularis (Gates and Mills 2005), hair cells of the cochlea (Lowell and Paparella 1977), and synapses between hair cells and spiral ganglion cells (Sergeyenko, Lall et al. 2013). Centrally, loss and shrinkage of spiral ganglion cells have been observed in post mortem studies of human patients with and without ARHL (Kirikae, Sato et al. 1964, Hansen and Reske-Nielsen 1965, Hinojosa and Nelson 2011). Similarly, pathologic changes throughout the central auditory system have been observed in deaf white cats (West and Harrison 1973, Schwartz and Higa 1982, Ryugo, Pongstaporn et al. 1997, Heid, Hartmann et al. 1998, Kral, Hartmann et al. 2002), and in the CN of congenitally deaf Shaker2-/- mice and DBA/2J mice with progressive high frequency hearing loss (Lee, Cahill et al. 2003, Connelly, Ryugo et al. 2017). It is therefore inferred that atrophy would occur in OC efferent neurons deprived of sensory stimulation.

Due to the reciprocal connectivity of the OC pathway to the auditory periphery, dysfunction to either IHCs or OHCs would cause the efferent system to malfunction. When IHC activation is disrupted in the case of conductive-, sensorineural- or genetic-hearing loss, there is reduced input to the central auditory system, resulting in less afferent input to MOC neurons. If degeneration occurs at the level of OHCs, MOC cells lose their peripheral target cells rendering them functionally ineffective. This situation has been reported in deaf hyperthyroid rats, where disordered OHC development results in abnormal axonal morphology of MOCs, with a 73% decrease in labelled fibers reaching the cochlea compared to normal rats (Cantos, Lopez et al. 2003). This observation suggests that OHCs are necessary for MOC axons to develop normally.

The changes in OC cell size and number observed here varied slightly from those reported previously, likely due to methodological differences. Cochlear ablation, a relatively severe manipulation, has previously been shown to result in a decrease in LOC efferent cell numbers observed at 2 weeks post-surgery, with this effect plateauing at around 2 months of age (Kraus and Illing 2005). In our material using more specific genetic ablations, efferent cell counts at 1 month of age were comparable across strains. Small changes in cell counts across OC efferent groups were observed in DBA/2J mice with progressive high frequency hearing loss at 3- and 6-months of age, but they were insignificant. Because these effects mirrored the age-related shifts observed in CBA/CaH mice, the changes may be explained by normal aging. In fact, the decrease in mean cell counts observed in the normal hearing animals from 3 to 6 months provides further evidence to an age-related component of OC efferent degeneration (Suthakar, Connelly et al. 2013, Radtke-Schuller, Seeler et al. 2015, Vicencio-Jimenez, Weinberg et al. 2021). As predicted, long-term deafness resulted in greater decreases in the size and number of OC efferents, reminiscent of that observed in cochlear ablation, albeit with a far longer time course.

We propose that the time course of changes to OC efferents observed in this study can be explained by the nature of peripheral pathology in our animals, which begins with failures of the stereocilia. Despite stereocilia malformation, hair cells remain in place for approximately 3 months in the congenitally deaf Shaker2-/- (Beyer, Odeh et al. 2000, Karolyi, Probst et al. 2003, Gong, Karolyi et al. 2006) and DBA/2J animals (Willott, Bross et al. 2005, Yang, Zhang et al. 2015). Additionally, while DPOAE thresholds were significantly elevated in both pathological hearing cases by 3 months of age, the persistence of measurable DPOAEs implies some residual OHC functionality, which is a considerably different situation compared to the cochlear-ablation and congenital-hypothyroidism conditions.

Cochlear ablation causes a generalized loss of both IHCs and OHCs across the entire frequency spectrum with unavoidable damage to the sensory neurons and local vasculature. On the other hand, the progressive hair cell loss observed in both DBA/2J (Yang, Zhang et al. 2015) and Shaker2-/- (Webster, Sobin et al. 1986) mice appears to be frequency specific, affecting basal (high frequency) cochlear regions first. Additionally, OHCs appear to be more vulnerable than IHCs in both DBA/2J (Hultcrantz and Spangberg 1997) and Shaker2-/- mice (Gong, Karolyi et al. 2006). Therefore, unlike the situation observed in hypothyroidism, because OHC pathology is limited to the stereocilia, we predicted that OC efferents in the congenitally deaf Shaker2-/-mouse would show normal axonal development. This hypothesis was verified by the observation that there was no significant difference observed in mean OC cell counts at 1 month of age. IHCs are retained in DBA/2J mice until at least 3 months of age, and OHC loss is specific to basal (high frequency) regions (Yang, Zhang et al. 2015). If the case of OC efferent degeneration is OHC loss, one would predict that given longer survival times, the situation in DBA/2J mice would start to resemble that seen in Shaker2-/- mice.

Hearing loss has a direct effect on the OC efferent system. Long-term congenital deafness results in a decrease in the number of contralaterally projecting cells, but only the size of ipsilateral efferents was affected. On the other hand, when some level of afferent input was retained in the case of specific high frequency hearing loss, there was no change in number of OC efferents but there were significant decreases in somatic size for contralateral MOC and ipsilateral LOC neurons. In adult chinchillas, aminoglycoside-induced hair cell loss causes peripheral efferent nerve fibres to degenerate with a faster time course relative to ANFs (McFadden, Ding et al. 2004). Additionally, cochleotomy in adult rats results in degeneration to OC efferents with a longer time course than seen in the CN, where progressive loss of intrinsic LOCs but not MOCs was observed for up to 3 months (Kraus and Illing 2005). Together, this evidence reveals that the time course of auditory efferent degeneration in the mature animal varies markedly from that observed in the afferent auditory system.

Previous studies employed immunohistochemical techniques to identify cholinergic OC neuron populations, the limitation of which is that subpopulations of MOC and LOC efferent neurons based on peripheral projection patterns are grouped together in subsequent analyses (Radtke-Schuller, Seeler et al. 2015, Vicencio-Jimenez, Weinberg et al. 2021). In this study, our use of neuronal tracing methods enabled characterization of contralateral vs ipsilateral populations. Unlike the case of cochlear ablation, which is typically unilateral, the underlying cochlear pathologies in the mutant mouse models used in this study are bilateral – like the situation in systemic models such as hypothyroidism and aminoglycoside-induced hearing loss. The differential effects observed in contralateral vs ipsilateral MOC neurons documented here provide clues regarding the underlying mechanisms resulting in the eventual loss of MOC neurons in our congenitally deaf model. Anatomically, afferent input to both populations of MOC neurons is symmetric and arises from the contralateral ear – IHCs synapse on type I ANFs which synapse on T-stellate neurons whose axons cross the midline to innervate MOC neurons (see Fig.2.8 of Brown 2011). The subpopulation distinction arises from the axonal projection pattern with respect to which ear MOC neurons innervate. Assuming that the afferent pathway is affected symmetrically when hearing dysfunction occurs bilaterally, the fact that contralateral MOC counts are more affected implies that axon length could be a significant factor contributing to greater loss of contralateral MOCs in long term congenital deafness.

In addition to primary afferent input from the cochlea, OC neurons also receive input from both descending auditory and neuromodulatory sources in the brain. In particular, both MOC and LOC neurons receive additional auditory input from the auditory cortex (Mulders and Robertson 2000, Horvath, Ribari et al. 2003, Brown, Mukerji et al. 2013, Hong, Zeppenfeld et al. 2022), the central nucleus of the inferior colliculus (Caicedo and Herbert 1993, Thompson and Thompson 1993, Malmierca, Le Beau et al. 1996, Suthakar and Ryugo 2017), and MNTB (Torres Cadenas, Fischl et al. 2020, Hong, Zeppenfeld et al. 2022). Histological studies suggest that both MOCs and LOCs appear to receive serotonergic and noradrenergic input (Woods and Azeredo 1999) and recent evidence confirms that serotonin mediated excitation of MOC neurons (Suthakar and Weisz 2023) contributes to protection from acoustic trauma (Ohata, Kondo et al. 2021). While inputs to OC neurons from within the auditory system will also lose primary afferent input from the cochlear in these mutant mouse strains, it is reasonable to speculate that the lack of gross changes to OC efferent morphology observed here may be due to plastic changes in neuromodulatory inputs arising from elsewhere in the brain. Further investigations into neuromodulation of OC efferent neurons are required.

In conclusion, our findings support previous reports that MOC but not LOC efferent neurons numbers decrease in normal aging. In young animals, MOC cell counts are comparable in models of hearing loss and deafness, but congenital deafness amplifies this age-related loss of MOC neurons significantly. Furthermore, subpopulations of MOC neurons display differential degeneration patterns with contralaterally projecting MOC neuron loss preceding the loss of ipsilaterally projecting MOC neurons. These data demonstrate that circuit level factors contributing to age-related MOC degeneration warrant more careful attention. It is unknown if MOC neuron output to the cochlear nucleus via axon collaterals are changed when they lose their primary peripheral axonal target. Such a situation would have important implications for brainstem feedback loops which could cause cascading plastic changes that significantly alter sound encoding in progressive hearing loss where some auditory function is retained. Further research is needed to understand the functional effects of changes to OC circuitry in age-related hearing loss.

## Acknowledgements

The authors gratefully acknowledge Katanyu Pongstaporn for his technical expertise in electron microscopy and Dr Michael Muniak who assisted with image analysis and provided valuable feedback during the drafting of this manuscript.

## Role of authors

All authors had full access to all the data in the study and take responsibility for the integrity of the data and the accuracy of the data analysis. Study concept and design: KS & DKR. Acquisition of data: KS & HDK. Analysis and interpretation of data: KS & DKR. Drafting of the manuscript: KS & DKR. Critical revision of the manuscript for important intellectual content: KS & DKR. Obtained funding: DKR. Administrative, technical, and material support: DKR. Study supervision: DKR.

## Abbreviations used

ARHL: Age related hearing loss
OC: Olivocochlear
MOC: Medial olivocochlear efferent
LOC: Lateral olivocochlear efferent
MEMR: Middle ear muscle reflex
SOC: Superior olivary complex
VNTB: Ventral nucleus of the trapezoid body
MNTB: Medial nucleus of the trapezoid body
LSO: Lateral Superior Olive
RPO: Rostral Periolivary Complex
DPOAE: Distortion product otoacoustic emissions
ABR: Auditory brainstem response
ChAT: Choline Acetyltransferase
FG: Fluorogold
CTB: Cholera Toxin Subunit-B
ICP: Inferior cerebellar peduncle
IHC: Inner Hair Cell
OHC: Outer Hair Cell

## Literature Cited

1. Albrecht, O., A. Dondzillo, F. Mayer, J. A. Thompson and A. Klug (2014). “Inhibitory projections from the ventral nucleus of the trapezoid body to the medial nucleus of the trapezoid body in the mouse.” Front Neural Circuits 8: 83.

2. Baashar, A., D. Robertson, N. J. Yates and W. Mulders (2019). “Targets of olivocochlear collaterals in cochlear nucleus of rat and guinea pig.” J Comp Neurol 527(14): 2273–2290.

3. Benson, T. E., D. K. Ryugo and J. W. Hinds (1984). “Effects of sensory deprivation on the developing mouse olfactory system: a light and electron microscopic, morphometric analysis.” J Neurosci 4(3): 638–653.

4. Beyer, L. A., H. Odeh, F. J. Probst, E. H. Lambert, D. F. Dolan, S. A. Camper, D. C. Kohrman and Y. Raphael (2000). “Hair cells in the inner ear of the pirouette and shaker 2 mutant mice.” J Neurocytol 29(4): 227–240.

5. Bogaerts, S., J. D. Clements, J. M. Sullivan and S. Oleskevich (2009). “Automated threshold detection for auditory brainstem responses: comparison with visual estimation in a stem cell transplantation study.” BMC Neuroscience 10(1): 1–7.

6. Borg, E. (1973). “On the neuronal organization of the acoustic middle ear reflex. A physiological and anatomical study.” Brain Res 49(1): 101–123.

7. Born, D. E. and E. W. Rubel (1985). “Afferent influences on brain stem auditory nuclei of the chicken: neuron number and size following cochlea removal.” J Comp Neurol 231(4): 435–445.

8. Brown, M. C. (1993). “Fiber pathways and branching patterns of biocytin-labeled olivocochlear neurons in the mouse brainstem.” J Comp Neurol 337(4): 600–613.

9. Brown, M. C. (2011). Anatomy of Olivocochlear Neurons. Auditory and Vestibular Efferents: 17–37.

10. Brown, M. C. and J. L. Levine (2008). “Dendrites of medial olivocochlear neurons in mouse.” Neuroscience 154(1): 147–159.

11. Brown, M. C., S. Mukerji, M. Drottar, A. M. Windsor and D. J. Lee (2013). “Identification of inputs to olivocochlear neurons using transneuronal labeling with pseudorabies virus (PRV).” J Assoc Res Otolaryngol 14(5): 703–717.

12. Caicedo, A. and H. Herbert (1993). “Topography of descending projections from the inferior colliculus to auditory brainstem nuclei in the rat.” J Comp Neurol 328(3): 377–392.

13. Campbell, J. P. and M. M. Henson (1988). “Olivocochlear neurons in the brainstem of the mouse.” Hear Res 35(2-3): 271–274.

14. Cantos, R., D. E. Lopez, J. A. Merchan and J. Rueda (2003). “Olivocochlear efferent innervation of the organ of corti in hypothyroid rats.” J Comp Neurol 459(4): 454–467.

15. Cantos, R., D. E. Lopez, M. L. Sala and J. Rueda (2000). “Study of the olivocochlear neurons using two different tracers, fast blue and cholera toxin, in hypothyroid rats.” Anat Embryol (Berl) 201(4): 245–257.

16. Conlee, J. W., T. N. Parks, C. Romero and D. J. Creel (1984). “Auditory brainstem anomalies in albino cats: II. Neuronal atrophy in the superior olive.” J Comp Neurol 225(1): 141–148.

17. Connelly, C. J., D. K. Ryugo and M. A. Muniak (2017). “The effect of progressive hearing loss on the morphology of endbulbs of Held and bushy cells.” Hear Res 343: 14–33.

18. Doucet, J. R. and D. K. Ryugo (2003). “Axonal pathways to the lateral superior olive labeled with biotinylated dextran amine injections in the dorsal cochlear nucleus of rats.” J Comp Neurol 461(4): 452–465.

19. Finch, C. E. (2003). “The biology of aging in model organisms.” Alzheimer Dis Assoc Disord 17 Suppl 2: S39–41.

20. Flood, D. G. and P. D. Coleman (1988). “Neuron numbers and sizes in aging brain: comparisons of human, monkey, and rodent data.” Neurobiol Aging 9(5-6): 453–463.

21. Frank, M. M., A. A. Sitko, K. Suthakar, L. Torres Cadenas, M. Hunt, M. C. Yuk, C. J. C. Weisz and L. V. Goodrich (2023). “Experience-dependent flexibility in a molecularly diverse central-to-peripheral auditory feedback system.” Elife 12.

22. Gates, G. A. and J. H. Mills (2005). “Presbycusis.” The Lancet 366(9491): 1111–1120.

23. Gong, T. W., I. J. Karolyi, J. Macdonald, L. Beyer, Y. Raphael, D. C. Kohrman, S. A. Camper and M. I. Lomax (2006). “Age-related changes in cochlear gene expression in normal and shaker 2 mice.” J Assoc Res Otolaryngol 7(3): 317–328.

24. Guinan, J. J. (2011). Physiology of the Medial and Lateral Olivocochlear Systems. Auditory and Vestibular Efferents. D. K. Ryugo and R. R. Fay, Springer New York. 38: 39–81.

25. Guinan, J. J., Jr., W. B. Warr and B. E. Norris (1984). “Topographic organization of the olivocochlear projections from the lateral and medial zones of the superior olivary complex.” J Comp Neurol 226(1): 21–27.

26. Hansen, C. C. and E. Reske-Nielsen (1965). “Pathological Studies in Presbycusis: Cochlear and Central Findings in 12 Aged Patients.” Arch Otolaryngol 82: 115–132.

27. Haug, H. (1987). “Brain sizes, surfaces, and neuronal sizes of the cortex cerebri: a stereological investigation of man and his variability and a comparison with some mammals (primates, whales, marsupials, insectivores, and one elephant).” Am J Anat 180(2): 126–142.

28. Heid, S., R. Hartmann and R. Klinke (1998). “A model for prelingual deafness, the congenitally deaf white cat--population statistics and degenerative changes.” Hear Res 115(1-2): 101–112.

29. Hinojosa, R. and E. G. Nelson (2011). “Cochlear nucleus neuron analysis in individuals with presbycusis.” The Laryngoscope 121(12): 2641–2648.

30. Hong, H., D. Zeppenfeld and L. O. Trussell (2022). “Electrical signaling in cochlear efferents is driven by an intrinsic neuronal oscillator.” Proc Natl Acad Sci U S A 119(44): e2209565119.

31. Horvath, M., O. Ribari, G. Repassy, I. E. Toth, Z. Boldogkoi and M. Palkovits (2003). “Intracochlear injection of pseudorabies virus labels descending auditory and monoaminerg projections to olivocochlear cells in guinea pig.” Eur J Neurosci 18(6): 1439–1447.

32. Hubel, D. H. and T. N. Wiesel (1970). “The period of susceptibility to the physiological effects of unilateral eye closure in kittens.” J Physiol 206(2): 419–436.

33. Hultcrantz, M. and M. L. Spangberg (1997). “Pathology of the cochlea following a spontaneous mutation in DBA/2 mice.” Acta Otolaryngol 117(5): 689–695.

34. Johnson, K. R., Q. Y. Zheng and L. C. Erway (2000). “A major gene affecting age-related hearing loss is common to at least ten inbred strains of mice.” Genomics 70(2): 171–180.

35. Kane, E. C. (1974b). “Patterns of degeneration in the caudal cochlear nucleus of the cat after cochlear ablation.” Anatomical Record 179: 67–92.

36. Karolyi, I. J., F. J. Probst, L. Beyer, H. Odeh, G. Dootz, K. B. Cha, D. M. Martin, K. B. Avraham, D. Kohrman, D. F. Dolan, Y. Raphael and S. A. Camper (2003). “Myo15 function is distinct from Myo6, Myo7a and pirouette genes in development of cochlear stereocilia.” Hum Mol Genet 12(21): 2797–2805.

37. Kirikae, I., T. Sato and T. Shitara (1964). “A Study of Hearing in Advanced Age.” Laryngoscope 74: 205–220.

38. Kirkpatrick, S., D. Ryugo and Z. Chaczko (2015). Development of a Web-Based System for Analysis of Auditory Brainstem Response (ABR) Data. Association for Research in Otolaryngology, Baltimore, Assoc. Res. Otolaryngol. Abs.

39. Kitzes, L. M. and M. N. Semple (1985). “Single-unit responses in the inferior colliculus: effects of neonatal unilateral cochlear ablation.” J Neurophysiol 53(6): 1483–1500.

40. Kral, A., R. Hartmann, J. Tillein, S. Heid and R. Klinke (2002). “Hearing after congenital deafness: central auditory plasticity and sensory deprivation.” Cereb Cortex 12(8): 797–807.

41. Kraus, K. S., D. Ding, H. Jiang, M. H. Kermany, S. Mitra and R. J. Salvi (2013). “Up-regulation of GAP-43 in the chinchilla ventral cochlear nucleus after carboplatin-induced hearing loss: correlations with inner hair cell loss and outer hair cell loss.” Hear Res 302: 74–82.

42. Kraus, K. S. and R. B. Illing (2004). “Superior olivary contributions to auditory system plasticity: medial but not lateral olivocochlear neurons are the source of cochleotomy-induced GAP-43 expression in the ventral cochlear nucleus.” J Comp Neurol 475(3): 374–390.

43. Kraus, K. S. and R. B. Illing (2005). “Cell death or survival: Molecular and connectional conditions for olivocochlear neurons after axotomy.” Neuroscience 134(2): 467–481.

44. Kreczmanski, P., H. Heinsen, V. Mantua, F. Woltersdorf, T. Masson, N. Ulfig, R. Schmidt-Kastner, H. Korr, H. W. Steinbusch, P. R. Hof and C. Schmitz (2007). “Volume, neuron density and total neuron number in five subcortical regions in schizophrenia.” Brain 130(Pt 3): 678–692.

45. Kuwabara, N. and J. M. Zook (1991). “Classification of the principal cells of the medial nucleus of the trapezoid body.” J Comp Neurol 314(4): 707–720.

46. Lee, D. J., H. B. Cahill and D. K. Ryugo (2003). “Effects of congenital deafness in the cochlear nuclei of Shaker-2 mice: an ultrastructural analysis of synapse morphology in the endbulbs of Held.” J Neurocytol 32(3): 229–243.

47. Li, D., C. M. Henley and B. W. O’Malley, Jr. (1999). “Distortion product otoacoustic emissions and outer hair cell defects in the hyt/hyt mutant mouse.” Hear Res 138(1-2): 65–72.

48. Liang, Y., A. Wang, I. A. Belyantseva, D. W. Anderson, F. J. Probst, T. D. Barber, W. Miller, J. W. Touchman, L. Jin, S. L. Sullivan, J. R. Sellers, S. A. Camper, R. V. Lloyd, B. Kachar, T. B. Friedman and R. A. Fridell (1999). “Characterization of the human and mouse unconventional myosin XV genes responsible for hereditary deafness DFNB3 and shaker 2.” Genomics 61(3): 243–258.

49. Lowell, S. H. and M. M. Paparella (1977). “Presbycusis: what is it?” Laryngoscope 87(10 Pt 1): 1710–1717.

50. Ma, Q., D. J. Anderson and B. Fritzsch (2000). “Neurogenin 1 null mutant ears develop fewer, morphologically normal hair cells in smaller sensory epithelia devoid of innervation.” J Assoc Res Otolaryngol 1(2): 129–143.

51. Malmierca, M. S., F. E. Le Beau and A. Rees (1996). “The topographical organization of descending projections from the central nucleus of the inferior colliculus in guinea pig.” Hear Res 93(1-2): 167–180.

52. McFadden, S. L., D. Ding, H. Jiang and R. J. Salvi (2004). “Time course of efferent fiber and spiral ganglion cell degeneration following complete hair cell loss in the chinchilla.” Brain Research 997(1): 40–51.

53. Moore, D. R. (1985). “Postnatal development of the mammalian central auditory system and the neural consequences of auditory deprivation.” Acta Otolaryngol Suppl 421: 19–30.

54. Mulders, W. H. and D. Robertson (2000). “Evidence for direct cortical innervation of medial olivocochlear neurones in rats.” Hear Res 144(1-2): 65–72.

55. Nordeen, K. W., H. P. Killackey and L. M. Kitzes (1983). “Ascending projections to the inferior colliculus following unilateral cochlear ablation in the neonatal gerbil, Meriones unguiculatus.” J Comp Neurol 214(2): 144–153.

56. Ohata, K., M. Kondo, Y. Ozono, Y. Hanada, T. Sato, H. Inohara and S. Shimada (2021). “Cochlear protection against noise exposure requires serotonin type 3A receptor via the medial olivocochlear system.” FASEB J 35(5): e21486.

57. Osen, K. K., E. Mugnaini, A. L. Dahl and A. H. Christiansen (1984). “Histochemical localization of acetylcholinesterase in the cochlear and superior olivary nuclei. A reappraisal with emphasis on the cochlear granule cell system.” Arch Ital Biol 122(3): 169–212.

58. Powell, T. P. and W. M. Cowan (1962). “An experimental study of the projection of the cochlea.” J Anat 96(Pt 2): 269–284.

59. Prieto, J. J., J. Rueda and J. A. Merchan (1990). “The effect of hypothyroidism on the development of the glycogen content of organ of Corti’s hair cells.” Brain Res Dev Brain Res 51(1): 138–141.

60. Probst, F. J., R. A. Fridell, Y. Raphael, T. L. Saunders, A. Wang, Y. Liang, R. J. Morell, J. W. Touchman, R. H. Lyons, K. Noben-Trauth, T. B. Friedman and S. A. Camper (1998). “Correction of deafness in shaker-2 mice by an unconventional myosin in a BAC transgene.” Science 280(5368): 1444–1447.

61. Radtke-Schuller, S., S. Seeler and B. Grothe (2015). “Restricted loss of olivocochlear but not vestibular efferent neurons in the senescent gerbil (Meriones unguiculatus).” Front Aging Neurosci 7: 4.

62. Roberts, B. L. and G. E. Meredith (1992). The Efferent Innervation of the Ear: Variations on an Enigma. The Evolutionary Biology of Hearing. D. B. Webster, A. N. Popper and R. R. Fay. New York, NY, Springer New York: 185–210.

63. Robertson, D., C. J. Anderson and K. S. Cole (1987). “Segregation of efferent projections to different turns of the guinea pig cochlea.” Hear Res 25(1): 69–76.

64. Ryugo, D. K., T. Pongstaporn, D. M. Huchton and J. K. Niparko (1997). “Ultrastructural analysis of primary endings in deaf white cats: morphologic alterations in endbulbs of Held.” J Comp Neurol 385(2): 230–244.

65. Schwartz, I. R. and J. F. Higa (1982). “Correlated studies of the ear and brainstem in the deaf white cat: changes in the spiral ganglion and the medial superior olivary nucleus.” Acta Otolaryngol 93(1-2): 9–18.

66. Sergeyenko, Y., K. Lall, M. C. Liberman and S. G. Kujawa (2013). “Age-related cochlear synaptopathy: an early-onset contributor to auditory functional decline.” J Neurosci 33(34): 13686–13694.

67. Sitko, A. A., M. M. Frank, G. E. Romero, M. Hunt and L. V. Goodrich (2025). “Lateral olivocochlear neurons modulate cochlear responses to noise exposure.” Proc Natl Acad Sci U S A 122(4): e2404558122.

68. Spangler, K. M., W. B. Warr and C. K. Henkel (1985). “The projections of principal cells of the medial nucleus of the trapezoid body in the cat.” J Comp Neurol 238(3): 249–262.

69. Spongr, V. P., D. G. Flood, R. D. Frisina and R. J. Salvi (1997). “Quantitative measures of hair cell loss in CBA and C57BL/6 mice throughout their life spans.” J Acoust Soc Am 101(6): 3546–3553.

70. Suthakar, K., C. J. Connelly and D. K. Ryugo (2013). Medial Olivocochlear Efferents: Changes in Shaker-2 Mice with Congenital Deafness. Association for Research in Otolaryngology 36th Annual Midwinter Meeting Baltimore, MD, Association for Research in Otolaryngology Abstracts 63.

71. Suthakar, K. and D. K. Ryugo (2017). “Descending projections from the inferior colliculus to medial olivocochlear efferents: Mice with normal hearing, early onset hearing loss, and congenital deafness.” Hear Res 343: 34–49.

72. Suthakar, K. and D. K. Ryugo (2021). “Projections from the ventral nucleus of the lateral lemniscus to the cochlea in the mouse.” J Comp Neurol 529(11): 2995–3012.

73. Suthakar, K. and C. Weisz (2023). “Serotonin excites medial olivocochlear neurons of the auditory efferent pathway.” The Journal of the Acoustical Society of America 154(4_supplement): A332–A332.

74. Thompson, A. M. and B. R. Schofield (2000). “Afferent projections of the superior olivary complex.” Microsc Res Tech 51(4): 330–354.

75. Thompson, A. M. and G. C. Thompson (1993). “Relationship of descending inferior colliculus projections to olivocochlear neurons.” Journal of Comparative Neurology 335(3): 402–412.

76. Torres Cadenas, L., H. Cheng and C. J. C. Weisz (2022). “Synaptic plasticity of inhibitory synapses onto medial olivocochlear efferent neurons.” J Physiol 600(11): 2747–2763.

77. Torres Cadenas, L., M. J. Fischl and C. J. C. Weisz (2020). “Synaptic Inhibition of Medial Olivocochlear Efferent Neurons by Neurons of the Medial Nucleus of the Trapezoid Body.” J Neurosci 40(3): 509–525.

78. Uziel, A., J. Gabrion, M. Ohresser and C. Legrand (1981). “Effects of hypothyroidism on the structural development of the organ of Corti in the rat.” Acta Otolaryngol 92(5-6): 469–480.

79. Van der Loos, H. and T. A. Woolsey (1973). “Somatosensory cortex: structural alterations following early injury to sense organs.” Science 179(4071): 395–398.

80. Vicencio-Jimenez, S., M. M. Weinberg, G. Bucci-Mansilla and A. M. Lauer (2021). “Olivocochlear Changes Associated With Aging Predominantly Affect the Medial Olivocochlear System.” Frontiers in Neuroscience Volume 15 - 2021.

81. Wang, Y. and P. B. Manis (2006). “Temporal coding by cochlear nucleus bushy cells in DBA/2J mice with early onset hearing loss.” J Assoc Res Otolaryngol 7(4): 412–424.

82. Warr, B. W. (1992). Organization of Olivocochlear Efferent Systems in Mammals. The Mammalian Auditory Pathway: Neuroanatomy. D. B. Webster, A. N. Popper and R. R. Fay. New York, Springer. 410–448.

83. Warr, W. B., J. B. Boche and S. T. Neely (1997). “Efferent innervation of the inner hair cell region: origins and terminations of two lateral olivocochlear systems.” Hear Res 108(1-2): 89–111.

84. Webster, D. B., A. Sobin and M. Anniko (1986). “Incomplete maturation of brainstem auditory nuclei in genetically induced early postnatal cochlear degeneration.” Acta Otolaryngol 101(5-6): 429–438.

85. West, C. D. and J. M. Harrison (1973). “Transneuronal cell atrophy in the congenitally deaf white cat.” J Comp Neurol 151(4): 377–398.

86. Willard, F. H. and D. K. Ryugo (1983). Anatomy of the central auditory system. The Auditory Psychobiology of the mouse. J. F. Willott. Springfield, IL, Charles C. Thomas: 201–304.

87. Willott, J. F., L. S. Bross and S. McFadden (2005). “Ameliorative effects of exposing DBA/2J mice to an augmented acoustic environment on histological changes in the cochlea and anteroventral cochlear nucleus.” J Assoc Res Otolaryngol 6(3): 234–243.

88. Willott, J. F., J. Kulig and T. Satterfield (1984). “The acoustic startle response in DBA/2 and C57BL/6 mice: relationship to auditory neuronal response properties and hearing impairment.” Hear Res 16(2): 161–167.

89. Woods, C. I. and W. J. Azeredo (1999). “Noradrenergic and serotonergic projections to the superior olive: potential for modulation of olivocochlear neurons.” Brain Res 836(1-2): 9–18.

90. Wu, J. S., E. Yi, M. Manca, H. Javaid, A. M. Lauer and E. Glowatzki (2020). “Sound exposure dynamically induces dopamine synthesis in cholinergic LOC efferents for feedback to auditory nerve fibers.” Elife 9.

91. Yang, L., H. Zhang, X. Han, X. Zhao, F. Hu, P. Li, G. Xie, L. Gao, L. Cheng, X. Song and F. Han (2015). “Attenuation of hearing loss in DBA/2J mice by anti-apoptotic treatment.” Hear Res 327: 109–116.

